# Co-activation of NF-κB and MYC renders cancer cells addicted to IL6 for survival and phenotypic stability

**DOI:** 10.1101/2020.04.12.038414

**Authors:** RR Barbosa, AQ Xu, D D’Andrea, F Copley, H Patel, P Chakravarty, A Clear, M Calaminici, M Janz, B Zhang, M Schmidt-Supprian, J Wang, JG Gribben, R Tooze, J Fitzgibbon, G Franzoso, K Rajewsky, DP Calado

## Abstract

NF-κB and MYC are found co-deregulated in human B and plasma-cell cancers. In physiology, NF-κB is necessary for terminal B-to-plasma cell differentiation, whereas MYC repression is required. It is thus unclear if NF-κB/MYC co-deregulation is developmentally compatible in carcinogenesis and/or impacts cancer cell differentiation state, possibly uncovering unique sensitivities. Using a mouse system to trace cell lineage and oncogene activation we found that NF-κB/MYC co-deregulation originated cancers with a plasmablast-like phenotype, alike human plasmablastic-lymphoma and was linked to t(8;14)[MYC-IGH] multiple myeloma. Notably, in contrast to NF-κB or MYC activation alone, co-deregulation rendered cells addicted to IL6 for survival and phenotypic stability. We propose that conflicting oncogene-driven differentiation pressures can be accommodated at a cost in poorly-differentiated cancers.

**Significance:** Our studies improve the understanding of cancer pathogenesis by demonstrating that co-deregulation of NF-κB and MYC synergize in forming a cancer with a poorly-differentiated state. The cancers in the mouse system share features with human Plasmablastic lymphoma that has a dismal prognosis and no standard of care, and with t(8;14)[MYC-IGH] Multiple myeloma, which is in overall resistant to standard therapy. Notably, we found that NF-κB and MYC co-deregulation uniquely render cells sensitive to IL6 deprivation, providing a road-map for patient selection. Because of the similarity of the cancers arising in the compound mutant mouse model with that of human Plasmablastic lymphoma and t(8;14)[MYC-IGH] Multiple myeloma, this model could serve in preclinical testing to investigate novel therapies for these hard-to-treat diseases.

**Highlights:**

- NF-κB and MYC co-activation originates (pre)plasmablast-like cancer
- NF-κB/MYC^+^ renders cancer cells addicted to IL6 for survival and phenotypic stability
- NF-κB/MYC^+^ cancers are alike a fraction of human plasmablastic lymphoma
- t(8;14)[MYC-IGH] multiple myeloma is linked to a NF-κB/MYC co-activation signature

## Introduction

Diffuse-large-B-cell-lymphoma (DLBCL) and Multiple myeloma (MM) are the most frequent hematological malignancies overall, each comprising multiple disease entities with different genetic profiles and response to treatment (Kumar et al., 2017; Young et al., 2019). There is no clear interconnection between these two diseases and reports of co-occurrence are extremely rare. However, DLBCL and MM share the same normal cell counterpart albeit at different stages of differentiation, i.e. a mature B-cell and a terminally-differentiated B-cell (Plasma-cell), respectively.

B-to-plasma cell differentiation is a multi-stage process that involves an intricate network of factors. It initiates through the downregulation of *PAX5* in an activated B-cell, a transcription factor critical for B-cell identity, allowing the expression of factors such as *XBP1* and *JCHAIN* (Nutt et al., 2015). Subsequent upregulation of *BLIMP1* and *IRF4* expression, at least in part downstream of the NF-κB pathway, is key for the reinforcement of the plasma-cell program and full terminal B-cell differentiation characterized by cell cycle arrest and substantial Ig secretion (Grumont and Gerondakis, 2000; Heise et al., 2014; Klein et al., 2006; Morgan et al., 2009; Nutt et al., 2015; Saito et al., 2007).

Notably, genetic alterations leading to the activation of the NF-κB pathway are found in ∼40% of DLBCLs corresponding primarily to the so-called activated B-cell subset (ABC-DLBCL) and in about 20% of MM patients (Annunziata et al., 2007; Compagno et al., 2009; Davis et al., 2010; Keats et al., 2007; Lenz et al., 2008a). However, in more than 80% of cases, MM cancer cells constitutively engage the NF-κB pathway through stimuli received from the cancer microenvironment (Demchenko and Kuehl, 2010; Hideshima et al., 2005; Staudt, 2010). NF-κB signaling plays a crucial role in the survival of mature B-cells and Plasma-cells in physiology and pathology and is critical for both ABC-DLBCL and MM cell lines (Staudt, 2010; Tornatore et al., 2014).

The knowledge that NF-κB directly induces the expression of genes essential in B-to-plasma cell differentiation and that ABC-DLBCL cancer cells despite being mature B-cells display features of plasmacytic differentiation suggest that a block of B-to-plasma cell is required in the pathogenesis of ABC-DLBCL. Consistently, the activity of BLIMP1, key for B-to-plasma cell differentiation, is lost exclusively in DLBCL of the ABC subtype through *BLIMP1* genetic aberrations (∼30% of cases), and indirectly by deregulated BCL6 expression from chromosomal translocations (∼26% of cases) (Mandelbaum et al., 2010; Pasqualucci et al., 2011; Tam et al., 2006; Zhang et al., 2015). We and others previously demonstrated using mouse models that disruption of *Blimp1* precluded B-to-Plasma cell differentiation and synergized with NF-κB activation for the development of lymphomas resembling ABC-DLBCL (Calado et al., 2010; Mandelbaum et al., 2010; Zhang et al., 2015).

The expression of the proto-oncogene MYC is deregulated through diverse mechanisms in most cancers, including ABC-DLBCL and MM (Anderson, 2011; Janz, 2006; Shaffer et al., 2006). In fact, around 70% of ABC-DLBCLs and at least 40% of MMs display MYC positivity at the protein level (Hu et al., 2013; Szabo et al., 2016; Xiao et al., 2014). Amongst other properties, MYC has a crucial role in regulating cell cycle entry of mammalian cells in both physiology and pathology and is critical in both ABC-DLBCL and MM cell lines(Holien et al., 2012; Lenz et al., 2008b; Shaffer et al., 2008).

The survival properties of the NF-κB pathway, and MYC’s role in cell cycle, favor the hypothesis that these factors are synergistic in carcinogenesis. However, it may be important in this hypothesis to account for specific oncogene-driven differentiation pressures. In fact, and in contrast to the key role of NF-κB in Plasma-cell differentiation it has been shown that MYC opposes this process and that *MYC* expression is repressed by BLIMP1 for terminal B-cell differentiation to ensue (Lin et al., 2000; Lin et al., 1997; Shaffer et al., 2002). It is thus unclear whether co-deregulation of NF-κB and MYC is developmentally compatible for carcinogenesis, and whether a potential synergy impacts the differentiation state of cancer cells, possibly uncovering unique sensitivities.

In this work we used a system in the mouse to trace cell lineage and oncogene activation and found that co-deregulation of NF-κB and MYC synergize to form cancer with a poorly-differentiated Plasma-cell state. The mouse cancers resembled a fraction of human Plasmablastic lymphoma, a rare disease with a dismal prognosis, and were linked at the gene expression level with MM carrying t(8;14)[MYC-IGH] that are in overall resistant to standard therapy(Montes-Moreno et al., 2010; Valera et al., 2010). In contrast to activation of either NF-κB or MYC alone, co-deregulation rendered cells sensitive to IL6 deprivation, and in the absence of IL6 the synergy between NF-κB and MYC in the formation of a cancer with a poorly-differentiated Plasma-cell state was lost. This work evidences that poorly-differentiated cancer cells can accommodate conflicting oncogene-driven differentiation pressures at the cost of a critical dependency.

## Results

### Experimental design

To investigate the oncogenic activity of constitutively active NF-κB signaling and MYC over-expression, alone or in combination, we used the *CD19*^*creERT2*^ transgene that targets Cre expression in B cells encompassing all stages of development except terminally differentiated B-cells (Plasma-cells) in a temporally regulated manner through tamoxifen administration (**Fig. 1**)(Yasuda et al., 2013). To induce activation of the NF-κB canonical pathway and/or MYC over-expression we generated compound mutant mice carrying the *CD19*^*creERT2*^ allele together with a *ROSA26* allele *IKK2ca*^*stopFL*^ and/or a *ROSA26* allele containing a *MYC* cDNA driven by a CAG promoter (**Fig. 1**A; *MYC*^*stopFL*^; (Calado et al., 2012; Sasaki et al., 2006)). Activation of the *IKK2ca*^*stopFL*^ allele by Cre-mediated recombination can be traced by expression of GFP, whereas activation of the *MYC*^*stopFL*^ allele is marked by expression of a signaling deficient truncated version of human CD2 (**Fig. 1A**). Mice carrying the *CD19*^*creERT2*^ transgene in combination with a *ROSA26* reporter allele containing a cDNA encoding YFP preceded by a *loxP* flanked STOP cassette (*YFP*^*stopFL*^; (Srinivas et al., 2001)) were used as controls (**Fig. 1B**).

**Figure 1.**
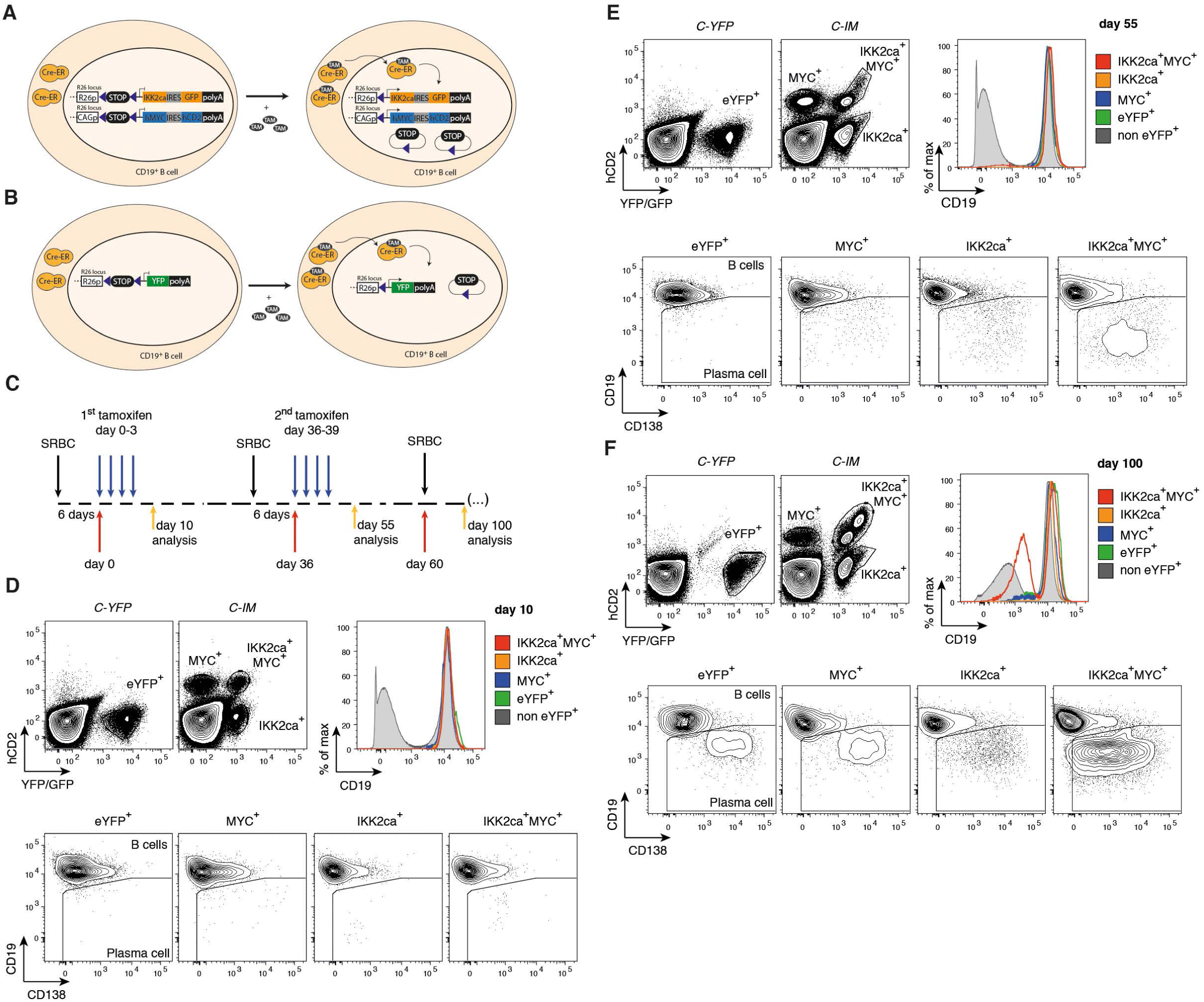
NF-κB signaling and MYC over-expression synergize for hyperplasia of a Plasma-cell like population. See also Fig S1. **(A-B)** Scheme illustrating the genetic systems used in the study. Triangles represent loxP sites. R26p and CAGp represent Rosa26 promoter and CAG promoter. **(B)** Schematic representation of the protocol of the study. Black arrow: immunization time-point with sheep red blood cells (SRBC); red arrow: day counting after the first tamoxifen administration; Blue arrow: tamoxifen administration time-point; yellow arrow: analysis time-point. **(D-F)** Representative flow cytometric analysis of Cre-mediated recombination in *C-YFP* and *C-IM* mice at day 10 (D), day 55 (E) and day 100 (F) after the first tamoxifen administration. Histograms represent CD19 expression within the recombined populations and bottom panels illustrate B-cell (CD19^+^) and Plasma-cell like (CD19^low^CD138^+^) populations within the individual reporter positive populations: GFP^neg^hCD2^+^ i.e. MYC^+^, GFP^+^hCD2^neg^ i.e. IKK2ca^+^, and GFP^+^hCD2^+^ i.e. IKK2ca^+^MYC^+^.

### *CD19cre*^*ERT2*^ allows the study of oncogenic mutations alone and in combination in a single mouse

In a *bona fide* model system of cancer the introduction of oncogenic mutations are tissue specific, temporally controlled, to trace the fate of cells in which mutations were introduced, and restricted to a small number of cells, mimicking the sporadic nature of oncogenic events. The ability to test the outcome of multiple oncogenes alone and in combination in the same mouse at the same time would be ideal to investigate synergy, dysergy or neutrality. We first immunized *CD19*^*creERT2*^ *IKK2ca*^*stopFL*^ *MYC*^*stopFL*^ (hereafter termed *C-IM*) and control *CD19*^*creERT2*^ *YFP*^*stopFL*^ (hereafter termed *C-YFP*) mice and injected tamoxifen at days 6, 7, 8, and 9 after immunization (**Fig. 1C**). Analysis at day 10 after the 1^st^ tamoxifen injection (i.e. at day 15 after immunization) revealed that control mice had a small fraction of YFP^+^ (∼2%) in the spleen (**Fig. 1D**). *C-IM* mice on other hand had three Cre-recombined populations: GFP^+^hCD2^neg^ cells (∼1%) representing cells with constitutive NF-κB activation alone, GFP^neg^hCD2^+^ cells (∼1%) representing cells where only MYC activation occurred, and GFP^+^hCD2^+^ (∼0.5%) representing cells carrying both mutations (**Fig. 1D**). The Cre-recombined populations in *C-YFP* and *C-IM* mice were overwhelmingly B-cells (**Fig. 1D**). We concluded that the *CD19*^*creERT2*^ system displays ideal properties to investigate the function of oncogenic events in carcinogenesis.

### NF-κB signaling and MYC over-expression synergize for hyperplasia of a Plasma-cell like population

To determine the impact of enforced NF-κB activation and MYC over-expression on B-cell fate, we established cohorts of control *C-YFP* mice, of experimental *C-IM* mice and mice carrying the *CD19*^*creERT2*^ allele in combination with either the *IKK2ca*^*stopFL*^ allele (hereafter termed *C-IKK2*) or *MYC*^*stopFL*^ (hereafter termed *C-MYC*) in accord with the experimental design in (**Fig. 1C**). We first examined the blood of mice for reporter positive cells at 55 days after the 1^st^ tamoxifen injection and multiple timepoints thereafter to trace their persistence or disappearance. The fractions of YFP^+^ cells in *C-YFP* and of MYC single expressing cells in *C-MYC* and *C-IM* decayed over the time of analysis, whereas the cellular population with enforced NF-κB activation in *C-IKK2* and *C-IM* remained constant (**Fig. S1A-C**). In contrast, the fraction of cells in which NF-κB and MYC were co-deregulated increased over the time of analysis (**Fig. S1A** and **B**). These data suggest that co-deregulation of NF-κB and MYC promote cellular expansion.

We next analyzed the spleens of mice at day 55 and 95 after tamoxifen injection and characterized phenotypically the reporter positive populations using flow-cytometry. Similarly to the analysis at day 10 (**Fig. S1D**) we found at these time-points a YFP^+^ population in *C-YFP* mice and three distinct reporter positive populations: GFP^neg^hCD2^+^ i.e. MYC^+^, GFP^+^hCD2^neg^ i.e. IKK2ca^+^, and GFP^+^hCD2^+^ i.e. IKK2ca^+^MYC^+^ (**Fig. 1E** and **F**). In contrast to the analysis at day 10, the IKK2ca^+^MYC^+^ population was at day 55 of analysis no longer homogenous with two distinct subpopulations emerging at day 95 (**Fig. 1E** and **F**). Further analysis using B-cell and Plasma-cell markers revealed the appearance of Plasma-cell like cells (CD19^low^CD138^+^), being particularly noticeable within IKK2ca^+^MYC^+^ cells (∼10% at day 55; ∼50% at day 95) compared to the other reporter positive cell populations (**Fig. 1E** and **F**). This data showed that expression of MYC from the *MYC*^*stopFL*^ allele did not impair the loss of the B-cell phenotype and acquisition of Plasma-cell like markers (Lin et al., 2000; Lin et al., 1997). Further, the data demonstrated that NF-κB signaling and MYC over-expression synergized for hyperplasia of a Plasma-cell like population.

### NF-κB or MYC expression leads to B cell lymphoma while co-deregulation to Plasma-cell like cancers

To assess a role of constitutive NF-κB signaling and MYC over-expression in carcinogenesis we aged control C-YFP, and experimental mice (*C-IKK2, C-MYC, C-IM*). *C-IM* displayed a dramatically reduced life span due to cancer occurrence (∼190 days after tamoxifen injection, p<0.0001) compared to all other genotypes (**Fig. 2A**). *C-IKK2* and *C-MYC* mice also succumbed to cancer to a variable degree, albeit at a much later time-point (∼500 days after tamoxifen injection; **Fig. 2A**). *C-MY*C and *C-IKK2* mice presented splenomegaly and accumulation of reporter positive cells in the spleen and lymph-nodes (**Fig. 2B**). In both cases, the cancer cells expressed the B-cell marker CD19 indicating B-cell lymphoma development (**Fig. 2C**). Macroscopic examination of cancer-bearing *C-IM* mice showed splenomegaly and hepatomegaly (**Fig. 2B**, not shown). Cancer-bearing *C-IM* mice displayed clonal accumulation of IKK2ca^+^MYC^+^ cells in the spleen, liver and bone marrow, with cells carrying single mutations being largely absent (**Fig. 2D** and **E** and **S2A**, not shown). Cancer cells in *C-IM* expressed the Plasma-cell marker CD138 (**Fig. 2D**), and histological examination of *C-IM* spleens showed compared to control mice a diffuse cell pattern with loss of follicular structure and B220 and Pax5 expression, but positivity for Irf4 and the proliferative marker Ki67 (**Fig. 2F**). Further suggesting a Plasma-cell like cancer, *C-IM* had an aberrant accumulation of IgM paraprotein (**Fig. S2B** and **C**). These data highlighted a unique synergy between NF-κB signaling and MYC over-expression for the formation of Plasma-cell like cancers.

**Figure 2.**
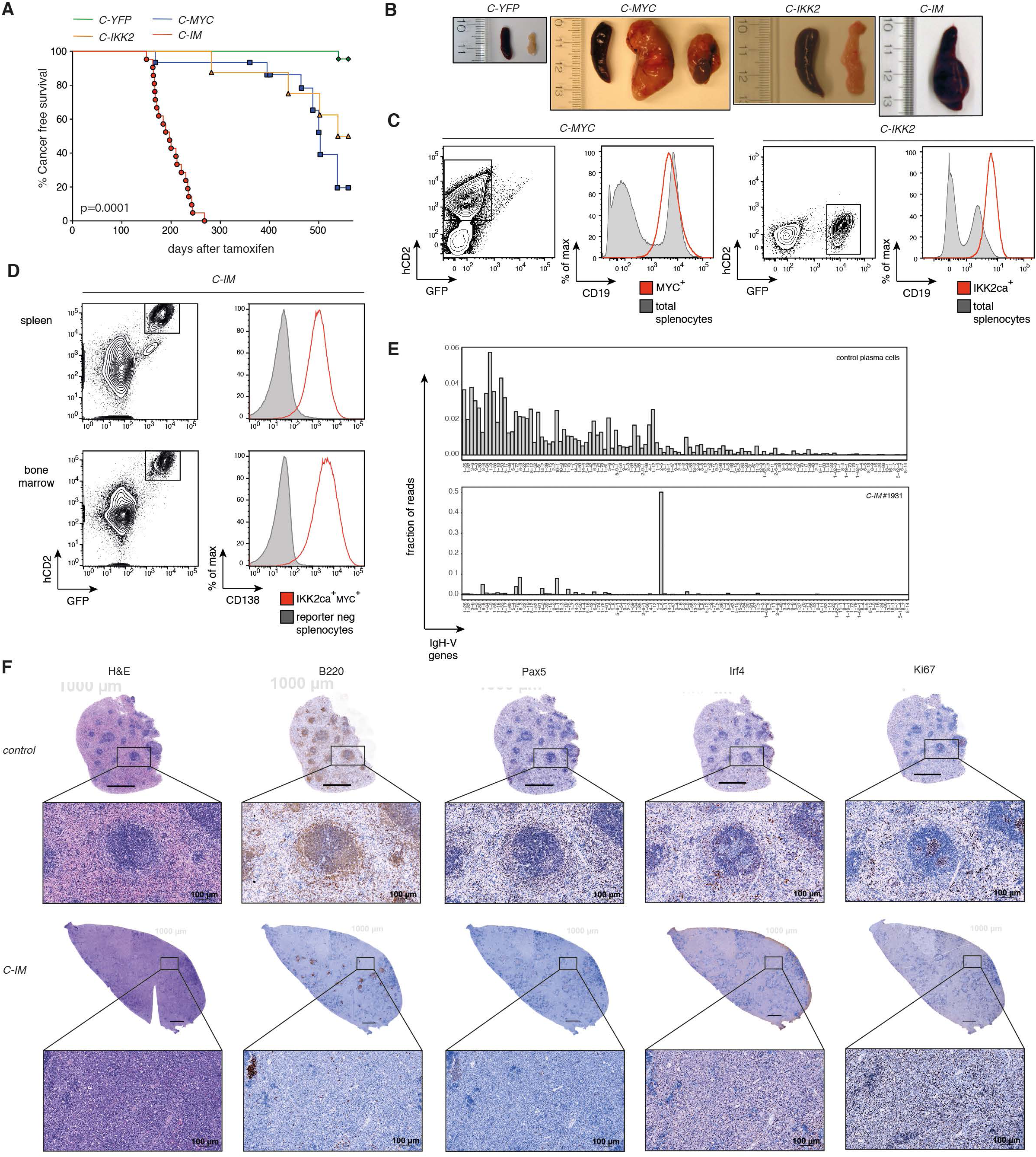
NF-κB or MYC expression leads to B cell lymphoma while co-deregulation to Plasma-cell like cancers. See also Fig S2. **(A)** Cancer free survival curve for control *C-YFP* mice, and experimental *C-IKK2, C-MYC*, and *C-IM* mice. **(B)** Representative images of spleen and mesenteric lymph nodes from aged mice of the indicated genotypes. **(C)** Representative flow cytometric analysis of cancers in spleen of *C-MYC* and *C-IKK2* mice. Left panels of each genotype show the frequency of reporter positive cells in the spleen; right panels of each genotype show the expression of CD19 within reporter positive cells (red), and total splenocytes from a control C57BL6 mouse (grey). **(D)** Representative flow cytometric analysis of cancers in spleen and bone marrow of *C-IM* mice. Left panels on each tissue show the frequency of double reporter positive cells; right panels on each tissue show the expression of the plasma cell marker CD138 within the double reporter positive population (red), and within the reporter negative population (grey). **(E)** Analysis of clonality by RNA sequencing. The fraction of reads mapped to each individual IgH-V gene out of all the reads mapped to IgH-V genes is shown. Top panel: control plasma cells (Shi et al., 2015), bottom panel: GFP^+^hCD2^+^ *C-IM* cancer cells (FACS purified). **(F)** Representative histological and immunohistochemical analysis of a *C-YFP* control spleen (top panels) and of a cancer in the spleen of *C-IM* mice (bottom panels) for H&E, B220, Pax5, Irf4, and Ki67.

### Cancers with NF-κB and MYC co-deregulation display a phenotype alike that of (pre)plasmablast

B-to-Plasma cell differentiation is a multi-stage process that involves an intricate network of factors (**Fig. 3A**, (Nutt et al., 2015)). To characterize the stage of B-to-Plasma cell differentiation of NF-κB^+^MYC^+^ cancer cells we performed gene expression profiling (GEP) by RNA sequencing of FACS-sorted GFP^+^hCD2^+^ cancer cells. We next compared the GEP of cancer cells with that of discrete B-cell and Plasma-cell populations (Shi et al., 2015). In agreement with the phenotypical characterization (**Fig. 2**), NF-κB^+^MYC^+^ cancer cells clustered with normal Plasma-cell populations in the loss of the expression of genes associated with the B-cell phenotype, including *Pax5, Ms4a1* (CD20), and *CD19* (**Fig. 3B**, and **C**). However, when analyzing genes which expression is increased in a Plasma-cell, the NF-κB^+^MYC^+^ cancer cells displayed an intermediate B-to-Plasma cell GEP, clustering on their own (**Fig. 3B**). Such an B-to-Plasma cell GEP state was highlighted by intermediate expression of *Blimp1, Irf4*, and *Xbp1* that are critical for B-to-Plasma cell differentiation and of other Plasma-cell expressed genes such as *CD138* (*Sdc1*) and *Jchain* (**Fig. 3B-D**). Gene signatures have previously been generated for Plasmablasts and Plasma-cells (Shi et al., 2015). Using these signatures we performed gene set enrichment analysis (GSEA) and found that the GEP of *C-IM* cancer cells was enriched for genes associated with Plasmablasts whereas the GEP of normal Plasma-cells was enriched for genes present in the Plasma-cell signature (**Fig. 3E**). These data suggested that enforced NF-κB activation and MYC over-expression synergized in the development of a cancer at a poorly-differentiated plasma-cell stage alike that of (pre)plasmablast.

**Figure 3.**
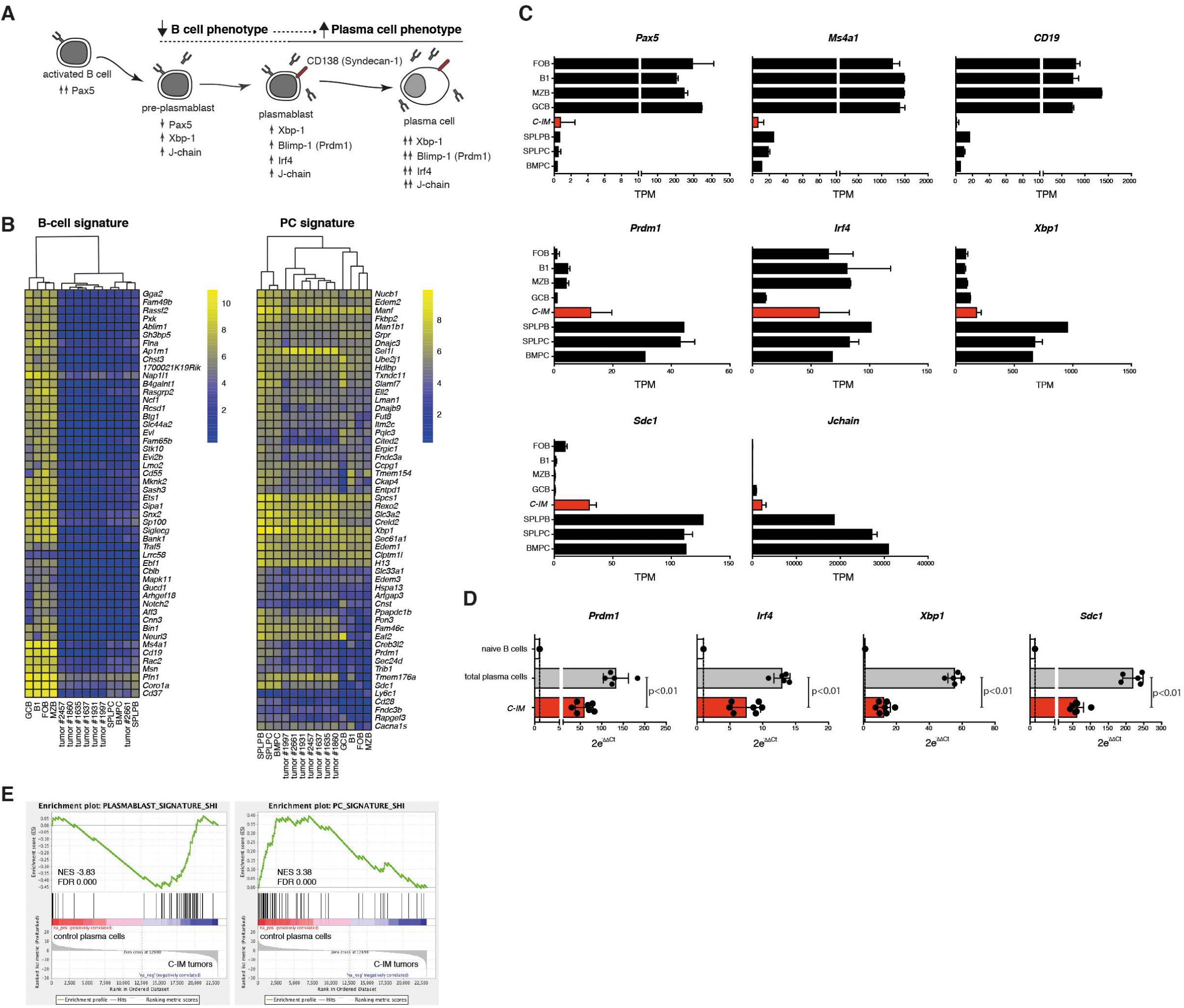
Cancers with NF-κB and MYC co-deregulation display a poorly-differentiated Plasma-cell state. **(A)** Schematic representation of the Plasma-cell differentiation process highlighting discrete populations and the associated expression of B-cell and Plasma-cell factors throughout the process. **(B)** Transcriptional analysis of *C-IM* cancers compared to discrete B-cell and Plasma-cell populations by RNAseq. GCB: Germinal Center B-cells, B1: B1 B-cells, FOB: Follicular B-cells, MZB: Marginal Zone B-cells, SPLPC: Spleen Plasma-cells, BMPC: Bone Marrow Plasma-cells, SPLPB: Spleen Plasmablasts (Shi et al., 2015). Seven *C-IM* cancers are depicted. B-cell signature: expression profile of the top 50 downregulated genes in BMPCs compared with FOBs, Plasma-cell signature: expression profile of the top 50 upregulated genes in BMPCs compared to FOBs, in addition to 4 genes of particular immunological interest (*Slc3a2, Prdm1, Ly6c1, Cd28*). Log2 FPKM expression values of genes are shown in the heatmaps, color-coded according to the legend. **(C)** RNAseq expression data for genes involved in B-cell identity and function (*Pax5, Ms4a1, CD19*) and factors related to Plasma-cell differentiation (*Prdm1, Irf4, Xpb1, Sdc1, Jchain*), in the populations mentioned in (B). TPM, transcripts per million. **(D)** Quantitative RT-PCR for plasma cell differentiation factors in seven *C-IM* cancers and spleen CD19^low^CD138^+^ plasma cells from C57BL6 mice. Data was normalized to a house-keeping gene (*Hprt1*) and then to the expression on naïve B cells (2e-ΔΔCt).

### NF-κB and MYC co-deregulation confers proliferative and survival advantage to (pre)Plasmablasts

To better understand the contribution of NF-κB and MYC co-deregulation we used a classical (pre)Plasmablast (B220^low^CD138^+^) differentiation assay *in vitro* in which B-cells are cultured in the presence of LPS (Andersson et al., 1972). For this purpose, we crossed the *IKK2ca*^*stopFL*^, *MYC*^*stopFL*^, and control *YFP*^*stopFL*^ alleles with *CD19*^*cre*^ that constitutively targets Cre expression in B-cells(Rickert et al., 1997). In agreement with the *in vivo* data, we found a profound synergy in the accumulation of B220^low^CD138^+^ cells in cultures derived from *CD19*^*cre*^ *IKK2ca*^*stopFL*^ *MYC*^*stopFL*^ B-cells compared to those where NF-κB or MYC deregulation occurred alone (**Fig. 4A** and **B**). Analysis of the fraction of B220^low^CD138^+^ cells per division revealed an increased proliferative capacity upon NF-κB and MYC co-deregulation compared to all other genotypes (**Fig. 4C)**. This increased proliferative capacity was accompanied by reduced apoptosis as measured by cleaved caspase 3 (**Fig. 4D**). These data showed that co-deregulation of NF-κB and MYC provided an advantage in proliferative capacity and survival of (pre)Plasmablasts.

**Figure 4.**
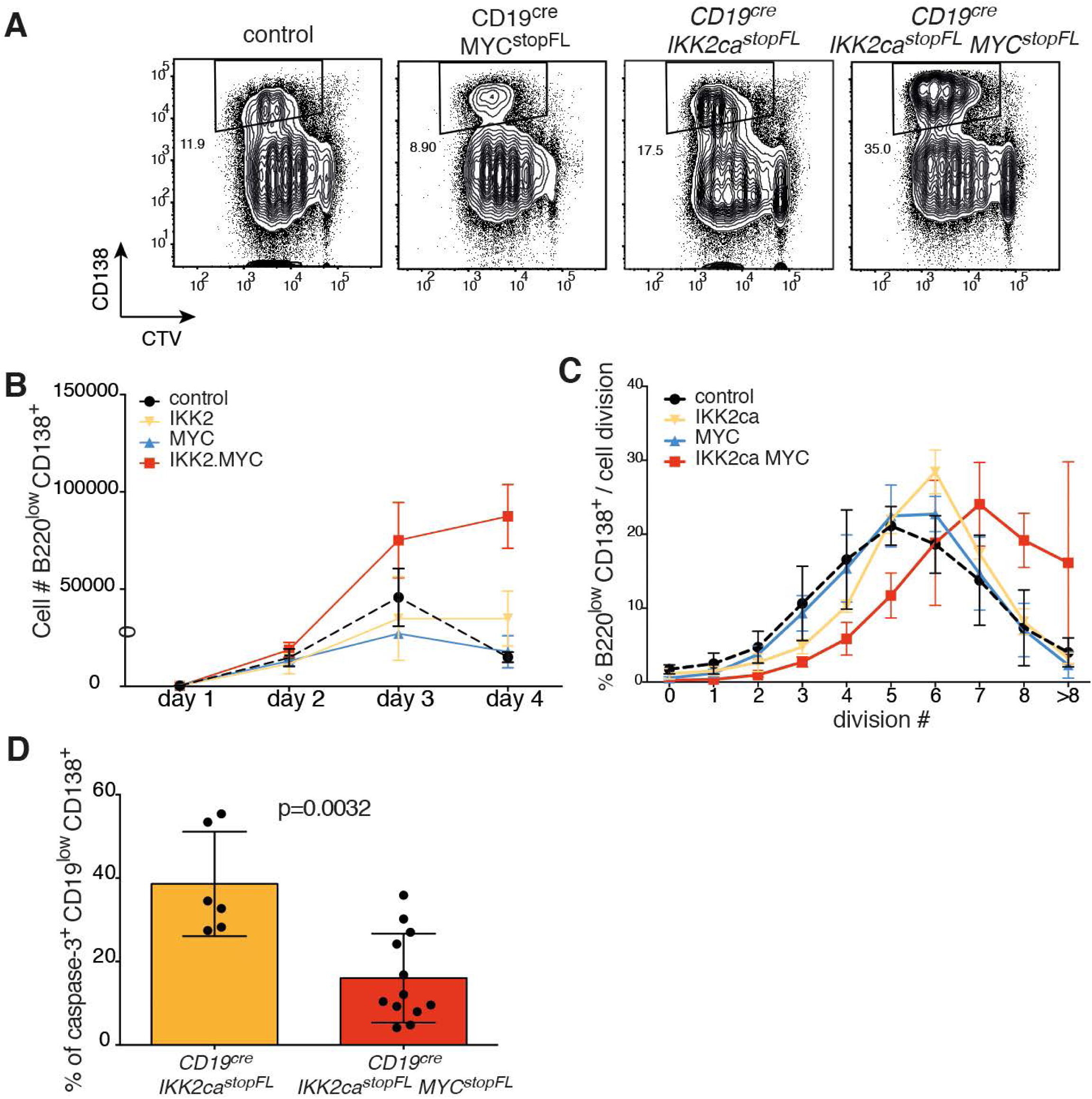
NF-κB and MYC co-deregulation confers proliferative and survival advantage to (pre)Plasmablasts. **(A)** Representative flow cytometric analysis of the proliferation profiles illustrating the frequency of (pre)plasmablasts (CD138^+^) on day 3 of LPS stimulation of B-cells from the indicated genotypes *in vitro*. Cell Trace Violet (CTV) dye dilution was used to assess proliferation. **(B)** Cell numbers of B220^low^CD138^+^ (pre)plasmablasts recovered from *in vitro* cultures at the indicated time-points. Black dashed lines represent control mice (control), yellow solid lines CD19cre IKK2ca^stopFL^ mice (IKK2ca), blue solid lines CD19cre MYC^stopFL^ (MYC), and red solid lines CD19cre IKK2ca^stopFL^ MYC^stopFL^ (IKK2caMYC). **(C)** Distribution of B220^low^CD138^+^ (pre)plasmablasts within each cell division as assessed by CTV dilution at day 4 of *in vitro* cultures. Mice genotypes are represented as in (B). **(D)** Frequency of cleaved caspase-3^+^ within CD19^low^CD138^+^ in spleen of mice assessed by *ex vivo* intracellular flow cytometric analysis. Mice of the indicated genotypes were analyzed between 16 and 29 weeks of age.

### NF-κB and MYC co-deregulation render cells addicted to IL6 for survival

To uncover dependencies of cancer cells with NF-κB and MYC co-deregulation we compared the GEP of the *C-IM* cancer cells with that of normal Plasma-cells. Compared to plasma-cells, the GEP of *C-IM* cancer cells was depleted for genes associated with the gene signature “Hallmarks_Apoptosis” **(Fig. 5A)** and pathway analysis revealed the enrichment of the “IL6 JAK/STAT” and “IL-6 signaling in MM” signatures in the GEP of *C-IM* cancer cells, reflected in part by the reduced expression of the STAT3 pathway inhibitor *Socs3*, and increased expression of *Il6st*, that encodes the IL6 co-receptor gp130 **(Fig. 5B-C)**. The IL6 JAK/STAT pathway was previously shown to play a key role in the survival of MM cancer cells *ex vivo* and *in vivo* (Klein et al., 1990a). IL6 ligation to IL6ra and gp130 leads to STAT3 phosphorylation (pSTAT3), homodimerization and nuclear translocation where it activates the transcription of multiple genes including the anti-apoptotic factors *BclxL* and *Mcl1* (Gaudette et al., 2014; Jourdan et al., 2003; Peperzak et al., 2013). Consistent with previous work demonstrating that IL6 is induced by NF-κB activation (Libermann and Baltimore, 1990), we found increased IL6 expression in the *in vitro* cultures derived from B-cells of mice carrying the *IKK2ca*^*stopFL*^ allele alone or in combination with *MYC*^*stopFL*^ (**Fig. S3A**). Notably, IL6 production was highly enriched in the (pre)Plasmablast population (∼85% of cells) compared to the activated B-cell population (∼15% of cells; **Fig. S3A**). To investigate the survival dependency on IL6 of (pre)Plasmablasts with NF-κB and MYC single or double de-regulation, we cultured B-cells with LPS in the absence or presence of an anti-IL6 neutralizing antibody (**Fig. 5D**). Suggestive of increased dependency and/or selection, the (pre)Plasmablasts with NF-κB and MYC co-deregulation showed the highest levels of pSTAT3 amongst all genotypes (**Fig. 5E**). Treatment of the cell cultures with anti-IL6 was effective in reducing the levels of STAT3 phosphorylation in all genotypes (**Fig. 5E**). The proliferative capacity of the (pre)Plasmablasts was unaltered by the anti-IL6 neutralizing antibody(**Fig. S3B**). However, we found a significant increase in the fraction of (pre)Plasmablasts marked for apoptosis in the condition where NF-κB and MYC were co-deregulated and a trend when MYC was deregulated alone (**Fig. 5F**). We next looked at the expression of anti-apoptotic proteins known to be downstream of IL6 signaling. With exception of the (pre)Plasmablasts carrying NF-κB activation alone, IL6 neutralization led to significant reduction in BclxL protein levels (**Fig. 5G**), whereas this was the case for Mcl1 protein levels in (pre)Plasmablasts carrying NF-κB/MYC co-deregulation and deregulation of MYC alone (**Fig. 5H**). These data showed that (pre)Plasmablasts with NF-κB and MYC co-deregulation and with deregulation of MYC alone are particularly sensitive to IL6 deprivation *in vitro*. Additional analysis by GEP demonstrated that such sensitivity to IL6 deprivation is (pre)Plasmablast specific, given the unaltered GEP profile of B cells with NF-κB/MYC co-deregulation upon IL6 neutralization (**Fig. 5I** and **J**). To investigate whether IL6 neutralization delayed cancer occurrence in *C-IM* mice, we aged cohort of mice and performed a single course of IL6 neutralization. Injection of anti-IL6 antibody significantly increased the length of cancer free survival of *C-IM* mice (**Fig. 5K**) suggesting a dependency of IKK2ca^+^MYC^+^ cancer cells on IL6 also *in vivo*.

**Figure 5.**
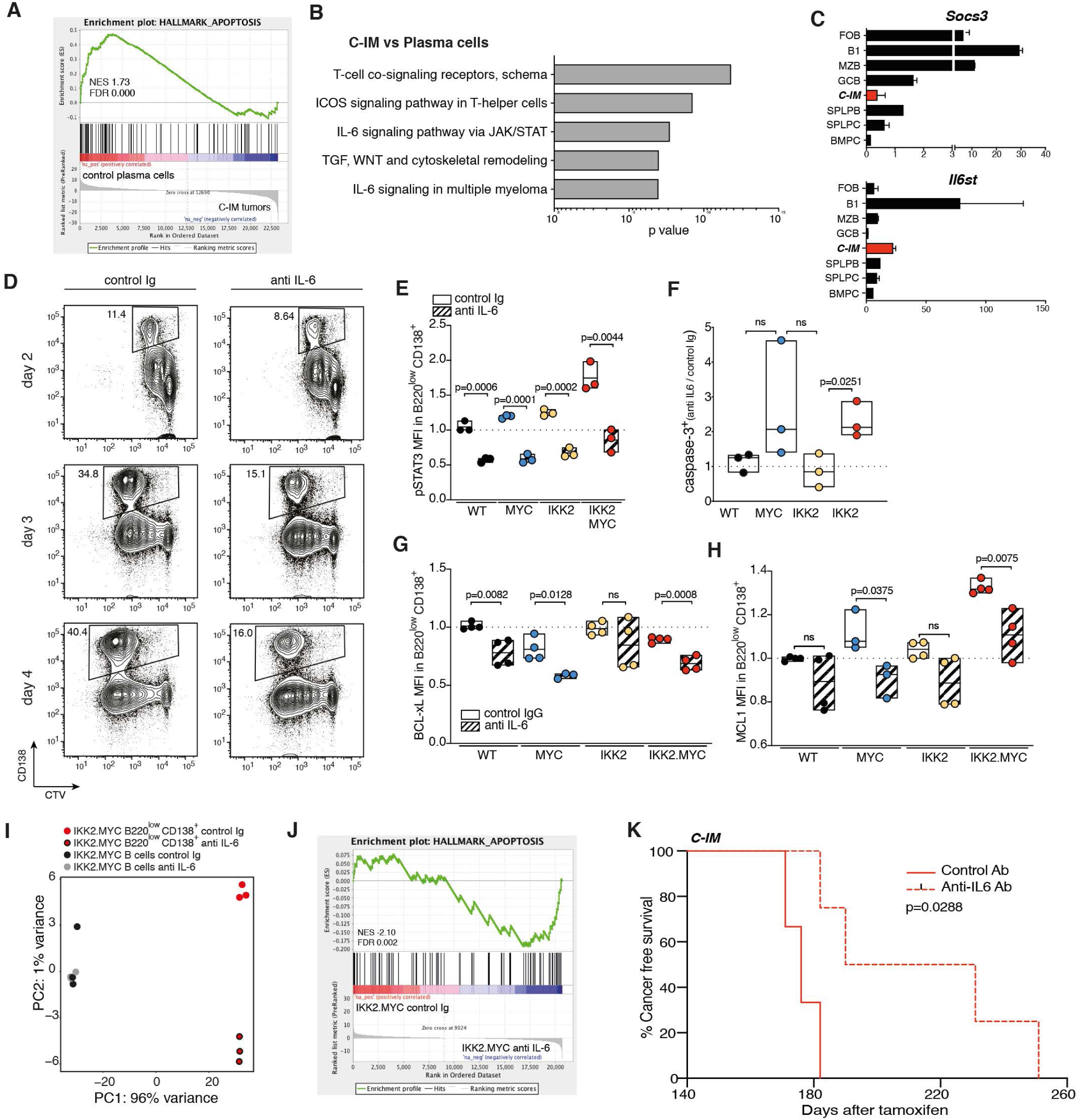
NF-κB and MYC co-deregulation render cells addicted to IL6 for survival. See also Fig S3. **(A)** Enrichment for genes in the signature “Hallmark_Apoptosis” using GSEA and the GEP of spleen CD19^low^CD138^+^ Plasma-cells from C57BL6 mice and *C-IM* cancer cells. **(B)** Pathway analysis enrichment (Metacore) of the GEP of *C-IM* cancer cells compared to spleen CD19^low^CD138^+^ Plasma-cells. The top 5 enriched pathways are shown. **(C)** RNAseq expression data for genes related to the IL-6 signaling pathway, *Socs3* and *Il6st*. GCB: Germinal Center B-cells, B1: B1 B-cells, FOB: Follicular B-cells, MZB: Marginal Zone B-cells, SPLPC: Spleen Plasma-cells, BMPC: Bone Marrow Plasma-cells, SPLPB: Spleen Plasmablasts (Shi et al., 2015), *C-IM*: GFP^+^hCD2^+^ *C-IM* cancer cells (FACS purified). **(D)** Representative flow cytometric analysis of the proliferation profiles illustrating the frequency of (pre)plasmablasts (CD138^+^) on day 2, 3, and 4 of LPS stimulation of B-cells from CD19cre IKK2ca^stopFL^ MYC^stopFL^ (IKK2caMYC) *in vitro*, in the presence of either control antibody (control Ig) or ant-IL6 neutralizing (anti-IL6). Cell Trace Violet (CTV) dye dilution was used to assess proliferation. **(E)** MFI of pSTAT3 in B220^low^CD138^+^ (pre)plasmablasts at day 2 of *in vitro* culture of B-cells from the indicated genotypes. Black circles represent control mice (WT), yellow circles CD19cre IKK2ca^stopFL^ mice (IKK2), blue circles CD19cre MYC^stopFL^ (MYC), and red circles CD19cre IKK2ca^stopFL^ MYC^stopFL^ (IKK2MYC). White background: control Ig, dashed background: anti-IL6. **(F)** Fold change (anti-IL6/control Ig) of the fraction of cleaved caspase 3^+^ cells within B220^low^CD138^+^ (pre)plasmablasts at day 2 of *in vitro* culture of B-cells from the indicated genotypes as in (E). **(G)** MFI of BCL-xL in B220^low^CD138^+^ (pre)plasmablasts at day 2 of *in vitro* culture of B-cells from the indicated genotypes as in (E). White background: control Ig, dashed background: anti-IL6. **(H)** MFI of MCL-1 in B220^low^CD138^+^ (pre)plasmablasts at day 2 of *in vitro* culture of B-cells from the indicated genotypes as in (E). White background: control Ig, dashed background: anti-IL6. **(I)** Principal component (PC) analysis plots of RNA-seq analysis of activated B cells (B220^+^) and B220^low^CD138^+^ (pre)plasmablasts at day 3 of *in vitro* culture of B-cells from CD19cre IKK2ca^stopFL^ MYC^stopFL^ (IKK2.MYC). **(J)** Enrichment for genes in the signature “Hallmark_Apoptosis” using GSEA and the GEP of B220^low^CD138^+^ (pre)plasmablasts at day 3 of *in vitro* culture of B-cells from CD19cre IKK2ca^stopFL^ MYC^stopFL^ in the presence of control Ig (IKK2.MYC control Ig) or ant-IL6 antibody (IKK2.MYC anti IL6). **(K)** Cancer free survival curve for *C-IM* mice treated with control Ig (solid red line) or ant-IL6 antibody (dashed red line).

### IL6 is critical for NF-κB^+^MYC^+^ (pre)plasmablast-like cancer phenotypic stability

We next wanted to determine whether IL6 dependency upon NF-κB and MYC co-deregulation was associated with cellular transformation at a poorly-differentiated Plasma-cell state. For that we generated compound mutant *C-IM* mice lacking IL6 (hereafter termed *C-IM-IL6KO*). Analysis of mice at 100 days after tamoxifen injection revealed a trend for a reduced fraction of cells with NF-κB and MYC co-deregulation, whereas cells with NF-κB or MYC deregulation alone showed either a trend for increased fraction of cells or no difference, respectively (**Fig. 6A** and **B**). When characterizing the fraction of CD19^low^CD138^+^ cells (Plasma-cell like) within each reporter positive population the impact of IL6 deprivation was unique to cells carrying NF-κB and MYC co-deregulation both in spleen (**Fig. 6C**) and bone marrow (**Fig. S4A**). Also only in the Plasma-cell like cells within the IKK2ca^+^MYC^+^ population we observed a significant increase in the fraction of cleaved caspase 3 positive cells (**Fig. 6D**). In contrast, IL6 deprivation had little impact on the cell cycle status of CD19^low^CD138^+^ cells within the IKK2ca^+^MYC^+^ population, whereas a slight increase in cells at the S/G2M phase was observed in the MYC activation alone condition (**Fig. 6E**). These data showed that NF-κB and MYC co-deregulation uniquely rendered cells sensitive to IL6 deprivation *in vivo*. These data also suggest that the apparent dependency of (pre)Plasmablasts with MYC deregulation alone on IL6 *in vitro* (**Fig. 5F** and **H**) could be a consequence of LPS induced NF-κB activation.

**Figure 6.**
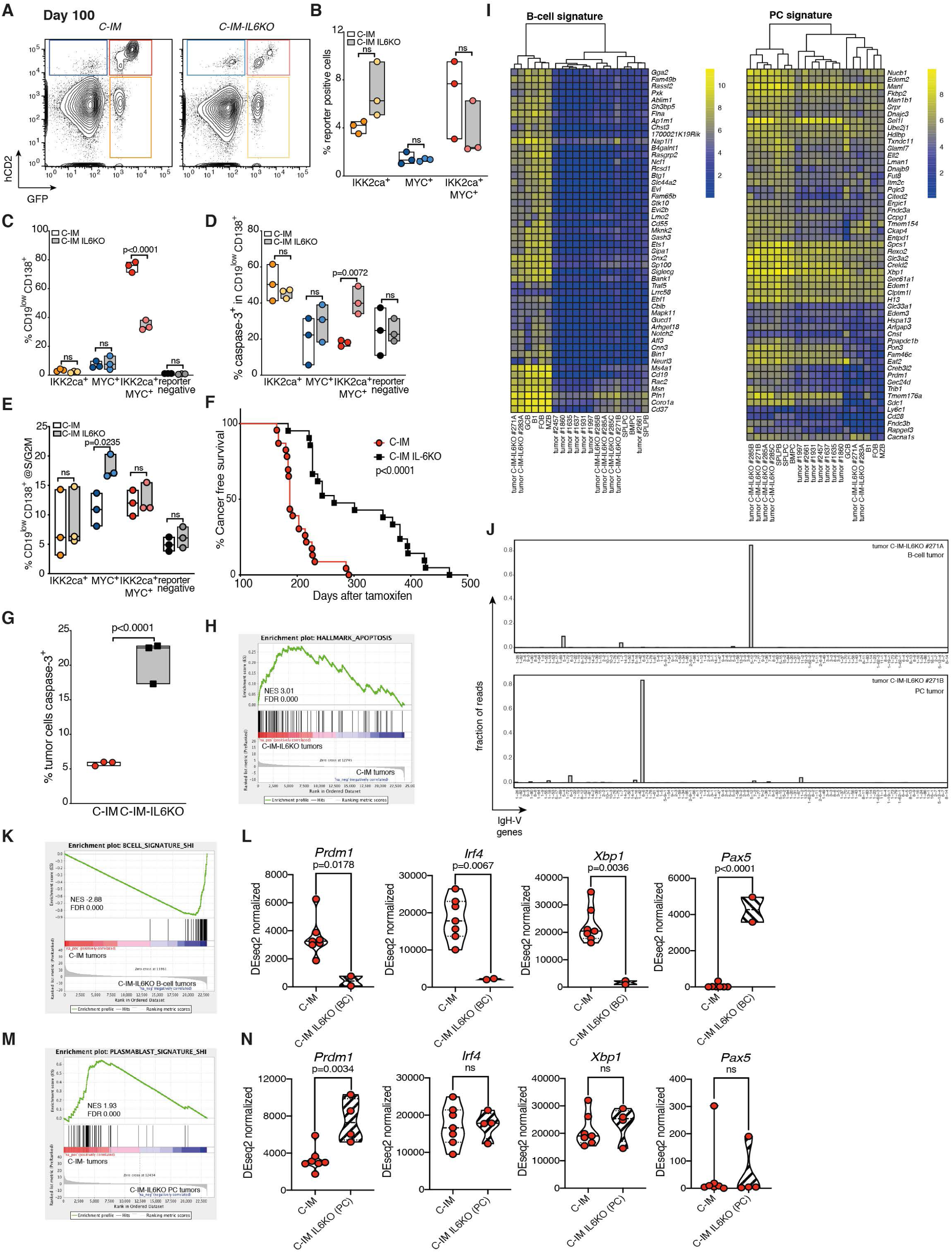
IL6 is critical for the formation of NF-κB^+^MYC^+^ cancers at a poorly-differentiated Plasma-cell state. See also Fig S4. **(A)** Representative flow cytometric analysis of Cre-mediated recombination in in spleen of *C-IM* and *C-IM-IL6KO* mice at day 100 after the first tamoxifen administration (protocol of study as in Fig. 1C). **(B)** Frequency in spleen of reporter positive populations: GFP^neg^hCD2^+^ i.e. MYC^+^, GFP^+^hCD2^neg^ i.e. IKK2ca^+^, and GFP^+^hCD2^+^ i.e. IKK2ca^+^MYC^+^. **(C)** Frequency in spleen of CD19^low^CD138^+^ cells within each reporter positive populations as in (B) and within reporter negative cells. **(D)** Frequency in spleen of cleaved caspase 3^+^ CD19^low^CD138^+^ cells within each reporter positive populations as in (B) and within reporter negative cells. **(E)** Frequency in spleen of CD19^low^CD138^+^ cells at the S/G2M phase of the cell cycle within each reporter positive populations as in (B) and within reporter negative cells. **(F)** Cancer free survival curve for *C-IM* and *C-IM-IL6KO* mice. **(G)** Frequency of cleaved caspase-3^+^ within CD19^low^CD138^+^ cancer cells of spleen of *C-IM* and *C-IM-IL6KO*. **(H)** Enrichment for genes in the signature “Hallmark_Apoptosis” using GSEA in the GEP of *C-IM-IL6KO* and *C-IM* cancer cells (GFP^+^hCD2^+^ FACS sorted). **(I)** Transcriptional analysis of *C-IM* and *C-IM-IL6KO* cancer cells compared to discrete B-cell and Plasma-cell populations by RNAseq. GCB: Germinal Center B-cells, B1: B1 B-cells, FOB: Follicular B-cells, MZB: Marginal Zone B-cells, SPLPC: Spleen Plasma-cells, BMPC: Bone Marrow Plasma-cells, SPLPB: Spleen Plasmablasts (Shi et al., 2015). Seven *C-IM* and six *C-IM-IL6KO* cancers are depicted. B-cell signature: expression profile of the top 50 downregulated genes in BMPCs compared with FoBs, Plasma-cell signature: expression profile of the top 50 upregulated genes in BMPCs compared to FOBs, in addition to 4 genes of particular immunological interest (*Slc3a2, Prdm1, Ly6c1, Cd28*). Log2 FPKM expression values of genes are shown in the heatmaps, color-coded according to the legend. **(J)** Analysis of clonality by RNA sequencing. The fraction of reads mapped to each individual IgH-V gene out of all the reads mapped to IgH-V genes is shown (Shi et al., 2015). Top panel: a GFP^+^hCD2^+^ *C-IM-IL6KO* B-cell cancer, bottom panel: a GFP^+^hCD2^+^ *C-IM-IL6KO* Plasma-cell cancer. **(K)** Enrichment for genes in the signature “Bcell_signature_Shi” using GSEA of the GEP of *C-IM* and *C-IM-IL6KO* (B-cell cancers) cancer cells. **(L)** RNAseq expression data for factors related to Plasma-cell differentiation (*Prdm1, Irf4, Xpb1*) and B-cell identity (*Pax5*) in cancer cells of *C-IM* and *C-IM-IL6KO* (B-cell cancers). **(M)** Enrichment for genes in the signature “Plasmablast_signature_Shi” using GSEA of the GEP of *C-IM* and *C-IM-IL6KO* (Plasma-cell cancers) cancer cells. **(N)** RNAseq expression data for factors related to Plasma-cell differentiation (*Prdm1, Irf4, Xpb1*) and B-cell identity (*Pax5*) in cancer cells of *C-IM* and *C-IM-IL6KO* (Plasma-cell cancers).

To investigate if IL6 had a role in the synergy between NF-κB and MYC in cell transformation we generated cancer cohorts of *C-IM* mice and *C-IM-IL6KO* mice following the previously described protocol of study (**Fig. 1B**). Compared to *C-IM* mice, *C-IM-IL6KO* had significantly prolonged cancer latency, with a median survival of 265 day for *C-IM-IL6KO* compared to 187 for *C-IM* (**Fig. 6F**). Notably, *C-IM-IL6KO* cancers had a significant increased fraction of cleaved caspase 3 positive cells compared to *C-IM* cancers and were enriched in their gene GEP for the gene signature “Hallmark_Apoptosis” (**Fig. 6G** and **H**). To better characterize the state of B-to-Plasma cell differentiation of *C-IM-IL6KO* cancer cells we performed GEP by RNA sequencing of FACS-sorted GFP^+^hCD2^+^ cancer cells and compared their GEP with that of discrete B-cell and Plasma-cell populations (Shi et al., 2015). Two out of 6 *C-IM-IL6KO* cancers analyzed by RNAseq clustered with the B-cell populations in the expression of genes associated with the B-cell phenotype, whereas 4/6 *C-IM-IL6KO* cancers clustered together with the *C-IM* cancers and the Plasma-cell populations in the loss of the expression of genes associated with the B-cell phenotype (**Fig. 6I**). Histological analysis revealed that roughly 15% (3/21) of *C-IM-IL6KO* cancers presented a B-cell like phenotype (**Fig. S4B**). When analyzing genes which expression is increased in Plasma-cells the four *C-IM-IL6KO* that for the B-cell signature clustered with *C-IM* no longer grouped together with these cancers (**Fig. 6I**). Whereas *C-IM-IL6KO* cancers clustered with the Plasma-cell populations (**Fig. 6I**), the *C-IM* cancers, as before (**Fig. 3B**) had an intermediate B-to-Plasma cell GEP, clustering on their own (**Fig. 6I**). Compared to *C-IM* cancers the *C-IM-IL6KO* B-cell like cancers had in a accordance to their phenotype reduced expression of *Blimp1, Irf4*, and *Xbp1*, whereas expression of *Pax5* was elevated (**Fig. 6K** and **L**). The *C-IM-IL6KO* Plasma-cell like cancers were depleted in genes characteristically expressed by Plasmablasts compared to *C-IM* cancers (**Fig. 6M**), and had increased expression of *Blimp1* (**Fig. 6N**). These data suggest that compared to the *C-IM* cancers, the *C-IM-IL6KO* Plasma-cell like cancers were more alike a well-differentiated Plasma-cell. In summary, IL6 was critical in the synergy between NF-κB and MYC deregulation for survival and transformation of a cell with a poorly-differentiated Plasma-cell phenotype. Phenotypic stability

### In the context of NF-κB activation MYC interferes with the expression of Plasma-cell factors

Previous work in cell lines suggested that repression of *MYC* by BLIMP1 is necessary for B cell terminal differentiation (Lin et al., 2000; Lin et al., 1997). In this work we showed that MYC over-expression driven from the *MYC*^*stopFL*^ allele does not interfere with the loss of the B-cell phenotype (**Fig. 3** and **4**). To better understand how MYC overexpression may interfere with the state of Plasma-cell differentiation we performed LPS stimulation of B-cells *in vitro* that either carrying NF-κB deregulation alone or together with MYC. We next purified reporter positive B220^low^CD138^+^ cells from these cultures and performed GEP using RNA-seq. Comparison of these GEP with previously derived Plasmablast and Plasma-cell signatures revealed that the GEP of cells with NF-κB and MYC co-deregulation was enriched for genes present in the Plasmablast signature whereas the GEP of cells with NF-κB deregulation alone enriched for genes present in the Plasma-cell signature (**Fig. 7A**). In agreement with these data cells with NF-κB deregulation alone expressed higher levels of genes key for the Plasma-cell differentiation process, namely *Blimp1, Irf4*, and *Xbp1* compared to cells with NF-κB and MYC co-deregulation (**Fig. 7B**). The level of the B-cell identity gene *Pax5* were identical between genotypes (**Fig. 7B**). These data indicates that MYC overexpression did not impact the loss of the B-cell phenotype, but it interfered with the reinforcement of the Plasma-cell program curtailing full Plasma-cell differentiation.

**Figure 7.**
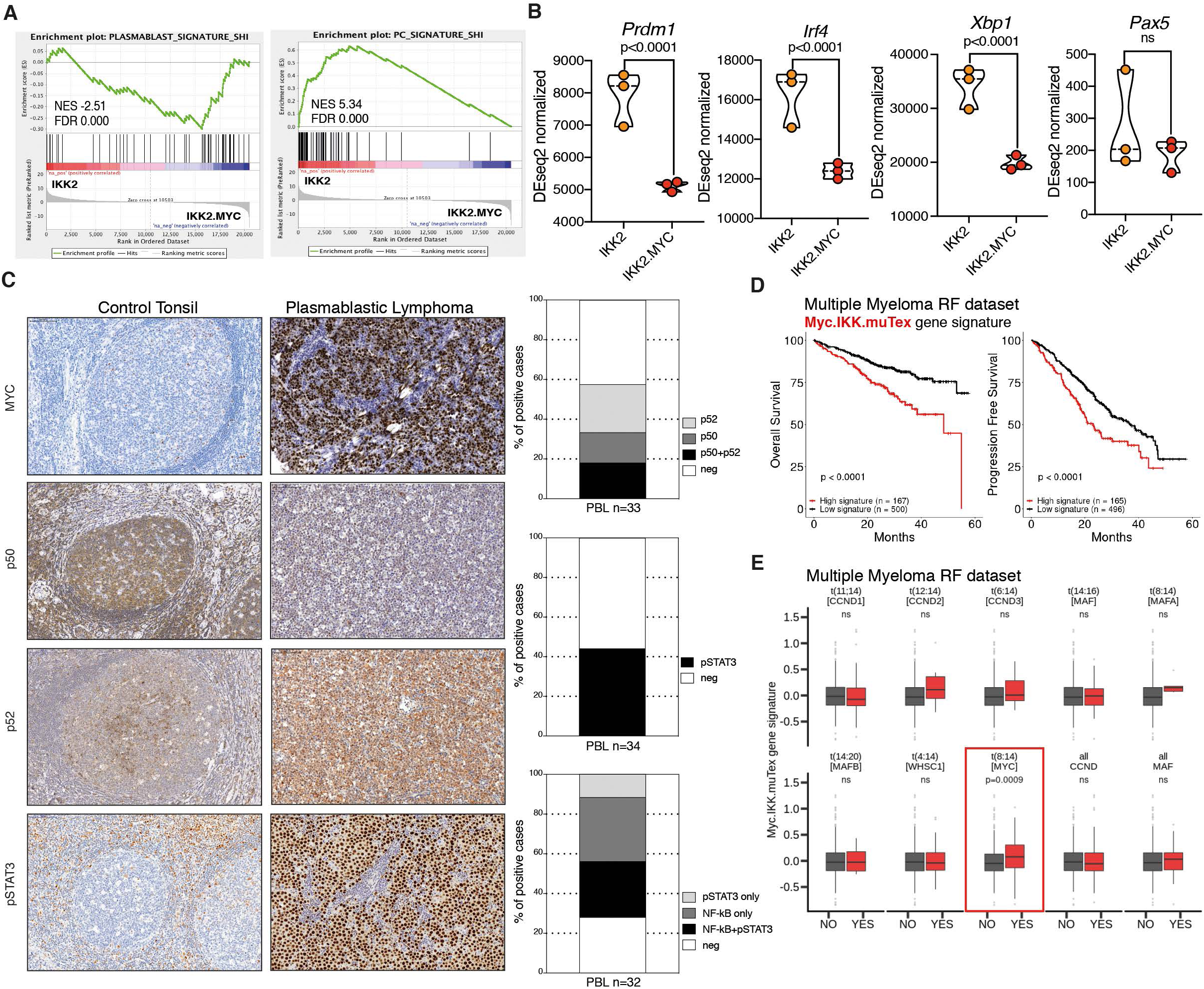
NF-κB/MYC^+^ cancers are alike a fraction of PBL and linked to t(8;14)[MYC-IGH] MM at the gene expression level. See also Fig S5. **(A)** Enrichment for genes in the signature “Plasmablast_signature_Shi” (left) and “PC_signature_Shi” (right) using GSEA of the GEP of B220^low^CD138^+^ (pre)Plasmablasts at day 3 of *in vitro* culture of B-cells from CD19cre IKK2ca^stopFL^ and CD19cre IKK2ca^stopFL^ MYC^stopFL^ mice. **(B)** RNAseq expression data for factors related to Plasma-cell differentiation (*Prdm1, Irf4, Xpb1*) and B-cell identity (*Pax5*) in B220^low^CD138^+^ (pre)Plasmablasts at day 3 of *in vitro* culture of B-cells from CD19cre IKK2ca^stopFL^ and CD19cre IKK2ca^stopFL^ MYC^stopFL^ mice. **(C)** Left, representative immunohistochemical analysis of human biopsies of tonsil and PBL for MYC, nuclear NF-κB p50, nuclear NF-κB p52, and phospho STAT3 (pSTAT3). A cutoff of >30% positive cells was used to determine positivity. Right, cumulative data of the immunohistochemical analysis. **(D)** Correlation between the gene enrichment of the “Myc.IKK2.muTex” signature with MM patient overall survival (left) and progression free survival (right; MMRF dataset). **(E)** Association between the “Myc.IKK2.muTex” signature and MM chromosomal abnormalities.

### *C-IM* cancers are alike a fraction of plasmablastic lymphoma

We next investigated if the cancers occurring in the *C-IM* mice resembled human disease. We excluded human-ABC-DLBCLs given that these lymphomas retain the B-cell phenotype in the vast majority of cells(Alizadeh et al., 2000). PBL on other hand is a human cancer that carries a Plasmablast-like phenotype and in which B-cell markers have been lost, despite being classified under DLBCL (Montes-Moreno et al., 2010; Swerdlow, 2008; Valera et al., 2010). Notably, at least 70% of PBL cases carry Ig/MYC translocations or *MYC* locus gain, and a recent PBL patient derived cell line was shown to be IL6 dependent (Mine et al., 2017; Taddesse-Heath et al., 2010; Valera et al., 2010). The *C-IM* cancers were generated through MYC overexpression, displayed a critical dependency on IL6 and had a poorly-differentiated Plasma-cell state similar to that of a (pre)Plasmablast. However, for the development of cancers in the mouse model system we also enforced the activation of the NF-κB pathway and currently it is unknown if PBL displays constitutive NF-κB signaling. We therefore assembled a cohort of 34 PBL cases and performed immunohistochemistry to determine MYC positivity, activation of the NF-κB canonical and alternative pathway through determination of nuclear p50 and p52, respectively, and activation of the STAT3 pathway using an anti-pSTAT3 antibody (Compagno et al., 2009). All of PBL cases analyzed were strongly positive for MYC, whereas ∼15% (5/33) were positive for nuclear p50, ∼25% (8/33) were positive for nuclear p52, and ∼15% (6/33) were positive for both, suggesting that ∼55% (19/33) of PBL cases in the study cohort display activation of the NF-κB pathway (**Fig 7C**). We also found that ∼44% (15/34) of PBL cases in the study cohort displayed phosphorylated STAT3 (**Fig 7C**). In summary, ∼28% (9/32) of PBL cases in the assembled cohort displayed co-deregulation of NF-κB, MYC and STAT3 phosphorylation (**Fig 7C**), indicating that the *C-IM* cancers are alike a fraction of PBL cancers.

### NF-κB and MYC co-deregulation identifies multiple myeloma patients with poor prognosis

We investigated whether oncogene associated gene signatures could also be predictive of patient outcome. Signatures associated with NF-κB or STAT3 activation were *per se* not predictive of patient outcome (**Fig. S5A** and **B**). In contrast, we found that signatures associated with MYC activity were highly predictive of patient outcome, in agreement to what was previously found at the protein level by others ((Moller et al., 2018); **Fig. S5C**). The lack of predictive value of both NF-κB or STAT3 gene signatures was puzzling. We therefore performed GEP of B220^low^CD138^+^ cells derived *in vitro* from B-cells without oncogene activation and with NF-κB and MYC deregulation alone or together. We next generated signatures of upregulated genes unique to each genotype i.e. specific to a condition where only enforced NF-κB occurred (IKK2.muTex), where only MYC de-regulation was induced (Myc.muTex), and for the condition where both NF-κB and MYC were enforced simultaneously (Myc.IKK2.muTex; **Fig S5D**). In contrast to the NF-κB and STAT3 gene signatures, all three muTex signatures were highly predictive of overall survival (OS) and progression free survival (PFS) (**Fig. 7D** and **S5E** and **F**). Notably, each muTex gene signature specifically enriched into discrete MM subsets according to their hallmark translocations (**Fig. 7E** and **S5E** and **F**). MM cases carrying either t(14;16)[IGH-MAF] or t(4;14)[FGFR3/WHSC1-IGH] positively correlated in their GEP with the IKK2.muTex signature whereas a negative correlation with this signature was observed for MM cases carrying t(8;16)[MYC-IGH] (**Fig. S5E**). The Myc.muTex gene signature was found to be significantly enriched in MM cases carrying t(6;14)[CCND3-IGH] (**Fig. S5F**), and the Myc.IKK2.muTex gene signature was significantly enriched in MM cases carrying t(8;16)[MYC-IGH] (**Fig. 7E**). These data suggest that gene signatures generated from phenotypically relevant cells with enforced NF-κB and MYC single and co-deregulation identify MM patients with poor prognosis and were linked to discrete genomic alterations providing a road-map for patient selection in specific therapeutic settings. The observation that a signature unique to NF-κB and MYC co-deregulation associates with MM carrying t(8;16)[MYC-IGH] indicates a role for the NF-κB and IL6 signaling pathway in these hard to treat MM patient subset.

## Discussion

In this work we studied oncogenic events that in physiology impose conflicting differentiation pressures. Specifically, we investigated carcinogenesis upon activation of NF-κB and MYC in the context of B-to-Plasma cell differentiation. We found that NF-κB and MYC co-deregulation synergized for the development of a cancer with a phenotype of a poorly-differentiated Plasma-cell, resembling a (pre)Plasmablast. Notably, in contrast to single NF-κB or MYC activation, co-deregulation rendered cells sensitive to IL6 deprivation, and IL6 was critical for the NF-κB/MYC synergy in forming a cancer with a poorly-differentiated Plasma-cell phenotype. We propose that poorly-differentiated cancer cells can accommodate at a cost conflicting oncogene-driven differentiation pressures.

The current work provides cues for the order and timing of mutation acquisition in the pathogenesis of mature B-cells. The finding that B-cell specific co-deregulation of NF-κB and MYC originates cancers devoid of B-cell markers supports that a block in B-to-Plasma cell differentiation, such as loss of BLIMP1 activity, is required for the pathogenesis of ABC-DLBCL. Given that cancer cells in ABC-DLBCL retain the B-cell phenotype, the cancers the current mouse model are not alike ABC-DLBCL. Instead, the NF-κB/MYC mouse cancers seem alike a fraction of human PBL and are linked with t(8;14)[MYC-IGH] MM at the gene expression level. We found that ∼55% of PBLs display NF-κB and MYC activation, and others have shown that a Plasmablastic morphology in MM is linked to high MYC expression and dismal survival prognosis (Lorsbach et al., 2011; Moller et al., 2018).

Surprisingly, *BLIMP1* genetic aberrations are also found in a fraction of PBL and MM (Chapman et al., 2011; Montes-Moreno et al., 2017). Their impact on BLIMP1 activity, timing of occurrence, disease formation and/or progression requires investigation. However, it is tempting to speculate that partial loss of BLIMP1 activity could alter the transcriptional landscape of cancer cells impacting the differentiation state and expression of cell cycle genes such as *MYC* (Montes-Moreno et al., 2017; Shaffer et al., 2008). Interestingly, *BLIMP1* genetic aberrations associate with poor prognosis in MM (Solimando et al., 2019).

Seminal work by Potter and colleagues described the development of Plasma-cell like cancers in mice upon pristane injection (Anderson and Potter, 1969). Additional studies revealed that the occurrence of such cancers involves chronic inflammation, IL6 signaling and the formation of *Myc* chromosomal translocations (Cheung et al., 2004; Dechow et al., 2014; Hilbert et al., 1995; Potter et al., 1985; Potter and Wiener, 1992; Rutsch et al., 2010; Suematsu et al., 1992). Other work showed that activation of MYC in B-cells to mostly originates cancers with a B-cell phenotype, although some mouse models displayed also Plasma-cell like cancers in a fraction of cases (Chesi et al., 2008; Harris et al., 1988; Kovalchuk et al., 2000; Park et al., 2005). It is accepted that NF-κB induces and is induced by inflammatory signals being considered the key link between inflammation and carcinogenesis (Taniguchi and Karin, 2018). However, despite this knowledge and that NF-κB favors B-to-Plasma cell differentiation, direct evidence of the participation of this pathway for the development of Plasma-cell like cancers was missing. Therefore, the current work provides a rationale for Plasma-cell like carcinogenesis in previous mouse models. The resemblance of the NF-κB/MYC mouse cancers with a fraction of PBLs and t(8;14)[MYC-IGH] MM indicates a role for the NF-κB pathway in the pathogenesis and/or progression of the cancers.

Our work and that of others suggests that knowledge on the differentiation state of cancer cells with respect to the normal cell counterpart allows a better definition of risk and of therapeutic opportunities (Paiva et al., 2017; Tarte et al., 2003). Human PBL and t(8;14)[MYC-IGH] MM patients have very poor prognosis and limited treatment options. The similarity of the NF-κB/MYC mouse cancers with these diseases may allow their use to identify therapeutic opportunities and strategies. We showed that a single course of anti-IL6 treatment was beneficial in delaying cancer progression. Supporting a possible role for IL6 in PBL, is the observation that this cytokine was required for the growth of a patient derived cell line (Mine et al., 2017). In MM, Inhibition of IL6 activity is of long-standing interest (Matthes et al., 2016). Initial proof of principle studies were highly promising, however, the results from randomized trials were unconvincing (Chari et al., 2013; Klein et al., 1991; Kurzrock et al., 2013; San-Miguel et al., 2014; Voorhees et al., 2013). It is possible that the results of randomized trials reflect existing MM cancer cell heterogeneity and the co-occurrence of IL6 dependent and independent clones (San-Miguel et al., 2014). This hypothesis is supported by work demonstrating that IL6 is not crucial in physiology for the survival of all Plasma-cells (Cassese et al., 2003). We found that B220^low^CD138^+^ cells carrying single MYC or NF-κB deregulation were IL6 independent *in vivo*. However, the failure of randomized trials and success of proof of principle studies could be at least in part due to patient selection (Klein et al., 1991; Klein et al., 1990b). Given this knowledge and the low toxicity of anti-IL6 treatment, although not curative, IL6/IL6R blocking may still be beneficial for a fraction of relapsed/refractory MM patients, particularly those displaying (pre)Plamablastic features and/or carrying t(8;14)[MYC-IGH].

## Supplemental Figure Legends

**Supplemental Figure 1.**
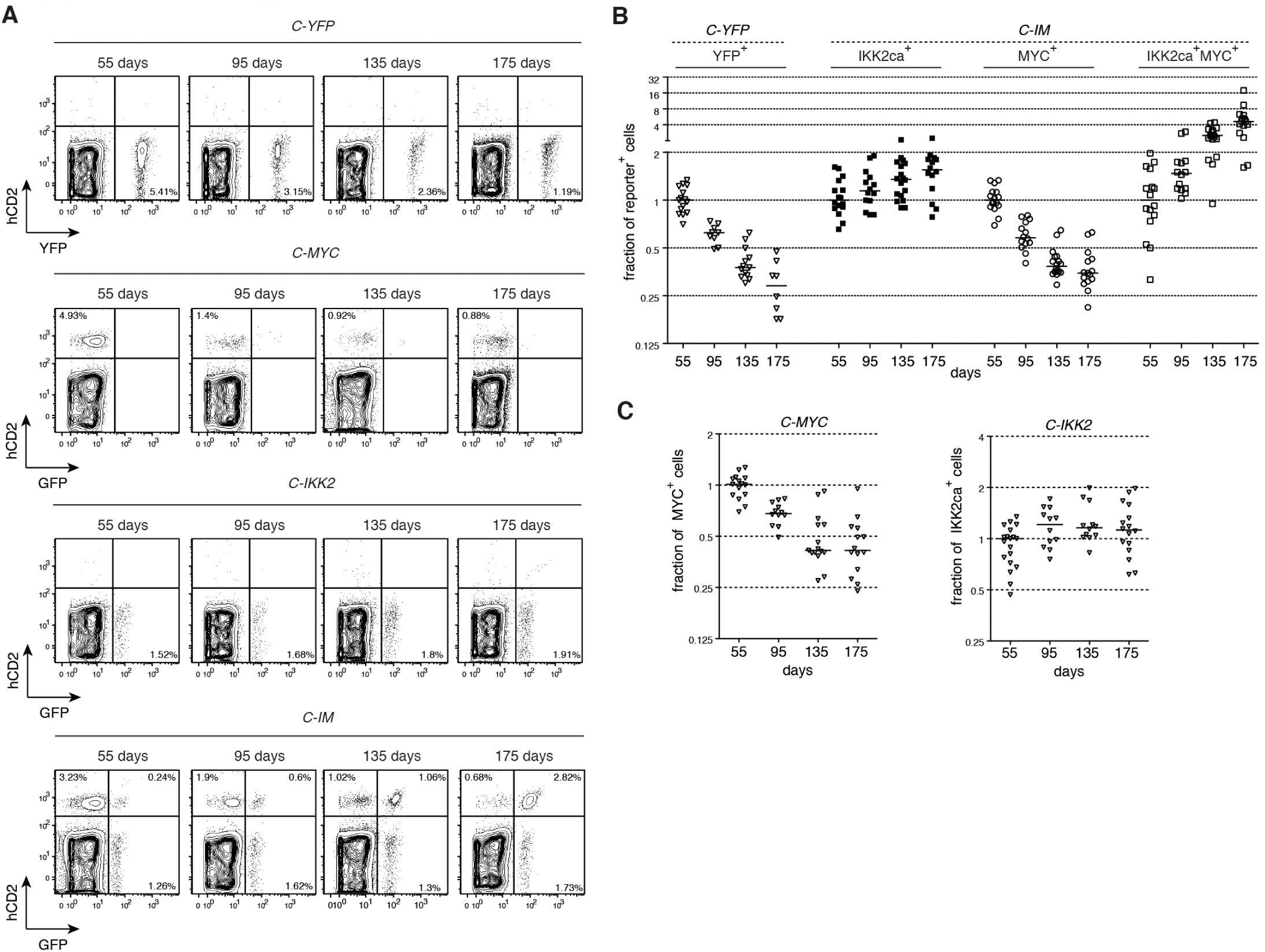
Dynamics of reporter positive subpopulations over-time. **(A)** Representative flow cytometric analysis of Cre-mediated recombination in the blood of *C-YFP, C-MYC, C-IKK2*, and *C-IM* at days 55, 95, 135, 175 after the first tamoxifen administration. **(B)** Fraction of cells within the individual reporter positive populations: YFP^+^ in *C-YFP* mice and GFP^neg^hCD2^+^ i.e. MYC^+^, GFP^+^hCD2^neg^ i.e. IKK2ca^+^, and GFP^+^hCD2^+^ i.e. IKK2ca^+^MYC^+^ in *C-IM* mice. Frequencies were normalized to day 55 after the first tamoxifen administration. **(C)** Left, fraction of GFP^neg^hCD2^+^ i.e. MYC^+^ in *C-MYC* mice. Right, fraction of GFP^+^hCD2^neg^ i.e. IKK2ca^+^ in *C-IKK2* mice. Frequencies were normalized to day 55 after the first tamoxifen administration.

**Supplemental Figure 2.**
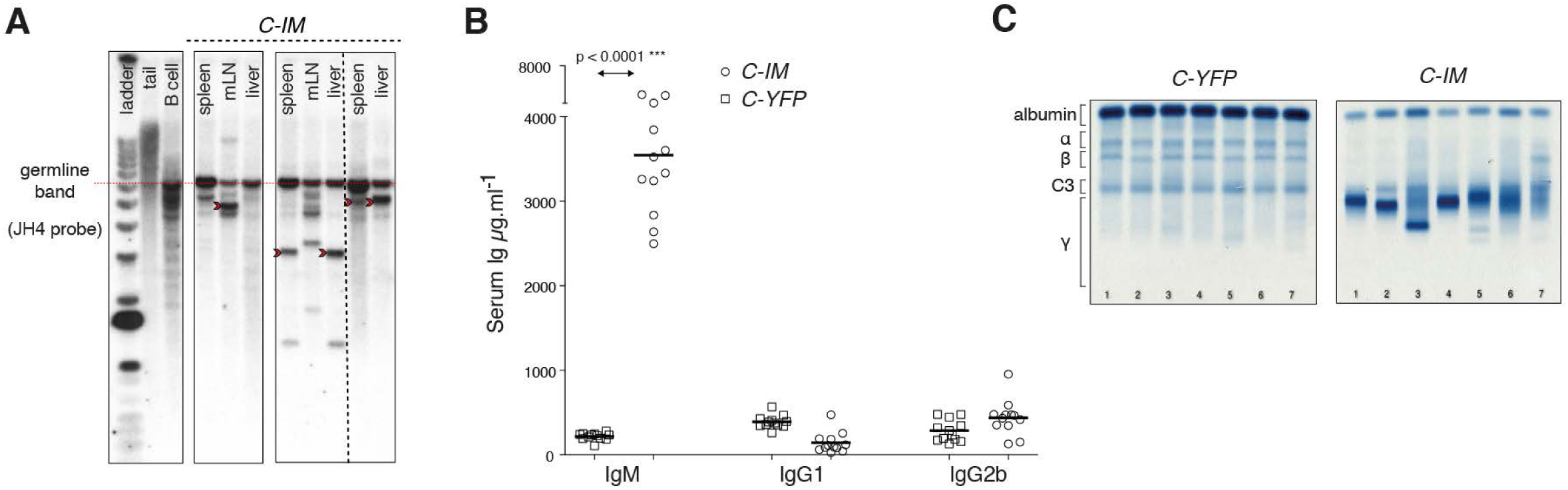
Characterization of *C-IM* cancer properties. **(A)** Southern blot analysis of cancer clonality using a JH4 probe. Dashed red line represents germline IgH configuration. Red arrow denotes clonal IgH. **(B)** ELISA of total serum IgM, IgG1 and IgG2b concentration in aged *C-YFP* and *C-IM* mice. **(C)** Serum protein electrophoresis of representative samples from aged *C-YFP* and *C-IM*. The position of albumin and of various globulin components of the serum is indicated.

**Supplemental Figure 3.**
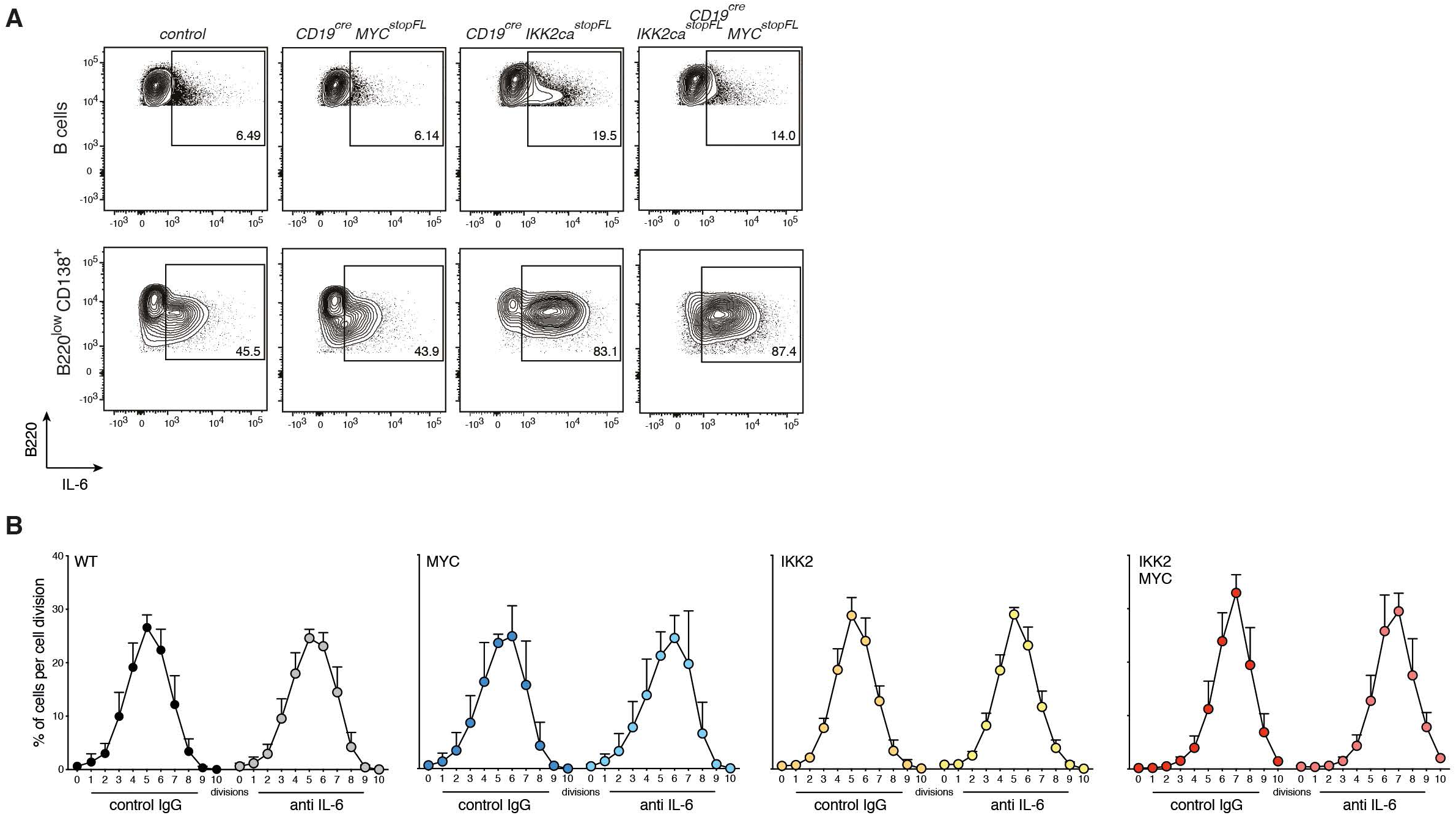
(pre)Plasmablasts are a source of IL6. **(A)** Representative flow cytometric analysis of intracellular staining for IL6 in *in vitro* day3 LPS stimulated B-cells from the indicated genotypes. Top panels: of activated B cells (B220^+^), Bottom panels: B220^low^CD138^+^ (pre)Plasmablasts. **(B)** Distribution of B220^low^CD138^+^ (pre)plasmablasts within each cell division as assessed by CTV dilution at day 4 of *in vitro* cultures in the presence of control Ig or anti-IL6 neutralizing antibody. Control mice (WT), CD19cre MYC^stopFL^ (MYC), CD19cre IKK2ca^stopFL^ mice (IKK2ca), CD19cre IKK2ca^stopFL^ MYC^stopFL^ (IKK2MYC).

**Supplemental Figure 4.**
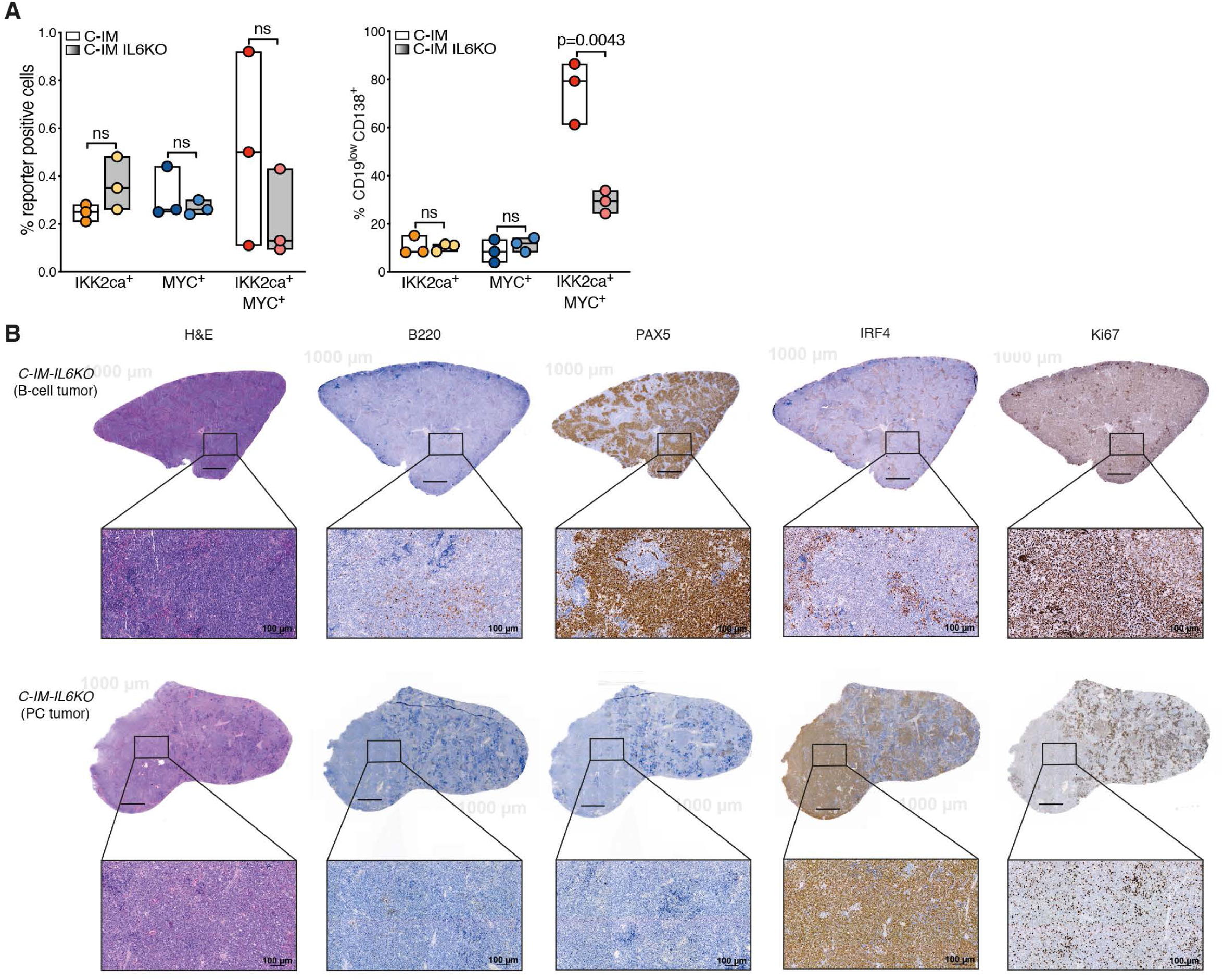
Analysis of *C-IM-IL6KO* mice. **(A)** Left, frequency in bone marrow of *C-IM* and *C-IM-IL6KO* mice at day 100 after first tamoxifen injection of reporter positive populations: GFP^neg^hCD2^+^ i.e. MYC^+^, GFP^+^hCD2^neg^ i.e. IKK2ca^+^, and GFP^+^hCD2^+^ i.e. IKK2ca^+^MYC^+^. Right, frequency in bone marrow of *C-IM* and *C-IM-IL6KO* mice at day 100 after first tamoxifen injection of CD19^low^CD138^+^ cells within each reporter positive populations and within reporter negative cells. **(B)** Representative histological and immunohistochemical analysis of a B-cell cancer in the spleen of a *C-IM-IL6KO* mice (top panels) and of a Plasma-cell cancer in the spleen of *C-IM-IL6KO* mice (bottom panels) for H&E, B220, Pax5, Irf4, and Ki67.

**Supplemental Figure 5.**
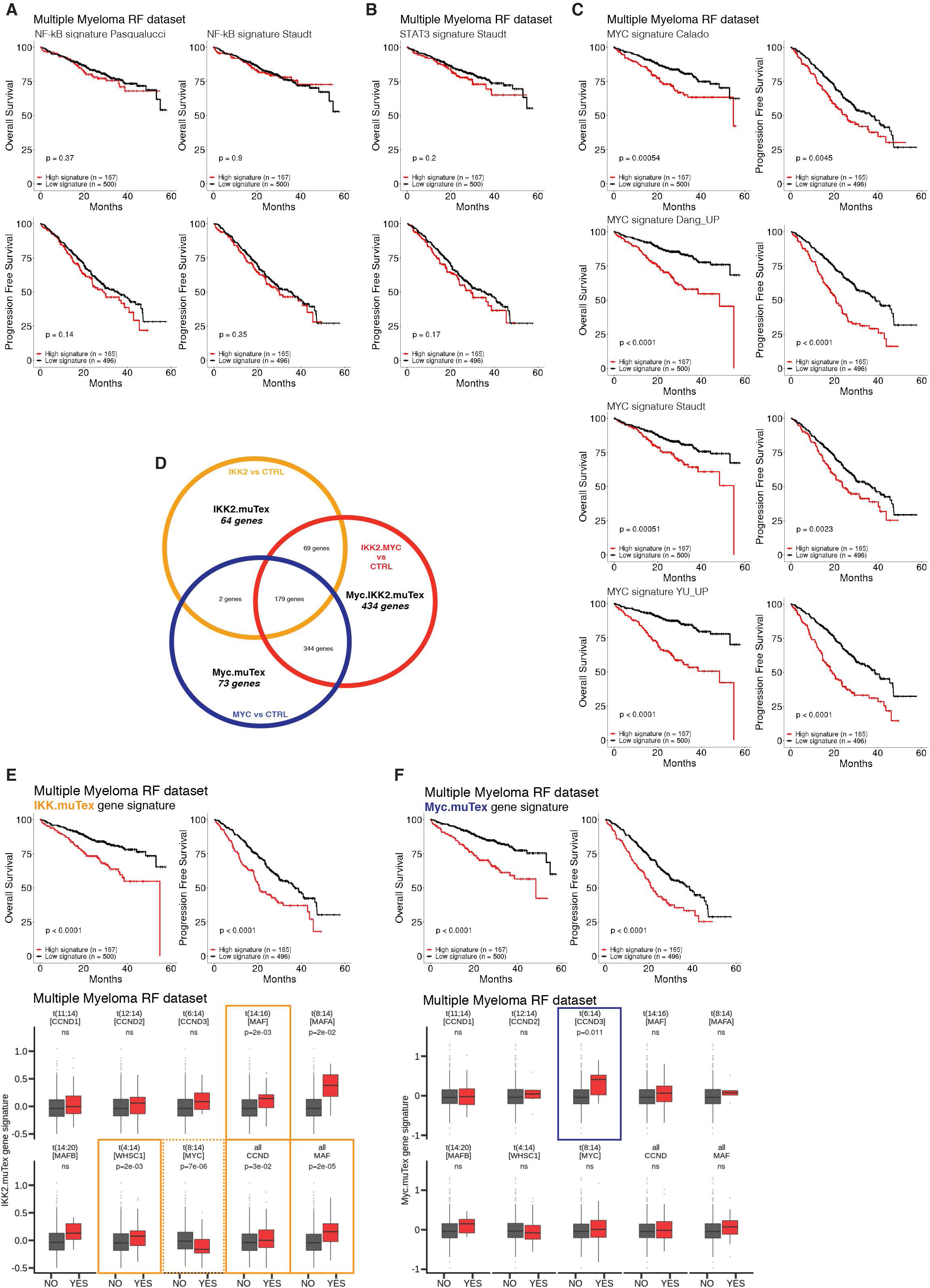
Gene signature enrichment in MM. **(A)** Correlation between the enrichment of a “NF-κB signature” (PASQUALUCCI) and the “NF-κB signature” (STAUDT) with MM patient overall survival (MMRF dataset). **(B)** Correlation between the enrichment of a “STAT3 signature” (STAUDT) with MM patient overall survival (MMRF dataset). **(C)** Correlation between the enrichment of multiple MYC signatures with MM patient overall survival (MMRF dataset). **(D)** Venn diagram showing the overlap in genes upregulated in the GEP of B220^low^CD138^+^ cells derived *in vitro* from B-cells of CD19cre IKK2ca^stopFL^ mice (IKK2), CD19cre MYC^stopFL^ (MYC), CD19cre IKK2ca^stopFL^ MYC^stopFL^ (IKK2MYC) mice. **(E)** Correlation between the enrichment of the “IKK2.muTex” signature with MM patient overall survival (left) and progression free survival (right; MMRF dataset). Association between the “IKK2.muTex” signature and MM chromosomal abnormalities. **(F)** Correlation between the enrichment of the “Myc.muTex” signature with MM patient overall survival (left) and progression free survival (right; MMRF dataset). Association between the “Myc.muTex” signature and MM chromosomal abnormalities.

## Materials and methods

### Mouse models and tumor cohorts

CD19^creERT2^, CD19^cre^, IKK2ca^stopFL^, MYC^stopFL^, IL6^KO/KO^, and YFP^stopFL^ alleles have been previously described. For T cell-dependent immunization, 8-to 12-week old mice were injected intravenously with 10^9^ defibrinated Sheep Red Blood Cells (SRBCs) (TCS Bioscience) in PBS. Mice were administered tamoxifen by oral gavage, dissolved in sunflower seed oil (both from Sigma). For full details on the experimental protocol, please see Figure 1E. Mouse cohorts were monitored twice a week for tumor development and euthanized if signs of tumor development occurred. Experiments were conducted using age-matched animals. Sex/gender was randomly distributed. All mice were on the C57BL/6 background. Mice were bred at The Francis Crick Institute under specific pathogen-free conditions. All animal experiments were carried out in accordance with national and institutional guidelines for animal care and were approved by The Francis Crick Institute Biological Resources Facility Strategic Oversight Committee (incorporating the Animal Welfare and Ethical Review Body) and by the Home Office, UK.

### In vivo treatment with anti-IL6 neutralizing antibody

To study the impact of IL6 on cancer progression, C-IM mice were treated with InVivoMAb anti-mouse IL-6 (bioXcell), or a control antibody (InVivoMAb rat IgG1 isotype control, anti-horseradish peroxidase; bioXcell). Administration of the antibodies was performed intraperitoneally three times per week for three weeks, starting on day 150 after tamoxifen administration. Mice were given 200ug of antibody per injection, diluted in InVivoPure dilution buffer, as per manufacturer’s instructions.

### In vitro stimulations

B cells were isolated from splenocyte suspensions using CD43 microbeads (Miltenyi Biotec), and plated at a density of 1×10^6^/mL in B-cell media (DMEM high glucose with Glutamax (Gibco), supplemented with Fetal Bovine Serum (GE Healthcare), non-essential aminoacids (Gibco), HEPES buffer (Gibco), sodium pyruvate (Gibco), Penicillin-Streptomycin (Gibco), and 2-Mercaptoethanol (Sigma-Aldrich)). B cells were stimulated with Lipopolysaccharide (LPS; Sigma-Aldrich) at a concentration of 10ug/mL. Where indicated, LPS stimulation was performed in the presence of blocking anti-IL-6 antibody or control IgG (Biolegend), at a concentration of 1ug/mL.

### Histology and Immunohistochemistry

Mouse spleens were fixed with 10% neutral buffered formalin (NBF) (Fischer Scientific) and embedded in paraffin. Sections were stained with hematoxylin and eosin (H&E) (Sigma). Antigen retrieval was performed with citrate buffer pH6, 23 minutes in the microwave. Ki67 staining was performed using the Ventana Discovery Ultra, CC1 for 48 minutes. All slides were counterstained with Harris Haematoxylin (Fisher Scientific), in Tissue Tek Prisma staining machine. Images were acquired with Zeiss Axio Scan.Z1 Slide Scanner and visualized with ZEN lite Blue Edition (Zeiss). For human Plasmablastic samples, tumor biopsies were fixed with 10% NBF and embedded in paraffin. Antigen retrieval (heat-induced epitope retrieval) was performed with citrate buffer using a pressure cooker and a commercial unmasking solution (Vector labs). The detection system used was from Biogenex (Super Sensitive Polymer HRP IHC Detection System). Images were acquired with the Pannoramic 250 Flash Scanner, uploaded into an online server (Casecenter) and visualized using Pannoramic Viewer (all 3DHistec).

### Flow cytometry

Single-cell suspensions of spleen and bone marrow were prepared in FACS buffer (2% FBS, 2 mM EDTA), in PBS (Gibco) and were treated with ACK Lysing Buffer (Gibco) for erythrocyte lysis. Single-cell suspensions were stained with antibodies, see Key Resources Table for details. Dead cells were excluded using Zombie NIR™ Fixable Viability Kit (BioLegend). For the assessment of cell division, cells were stained with CellTrace™ Violet (Invitrogen) as per manufacturer’s specifications, prior to in vitro stimulation. For the detection of cleaved caspase-3, MCL1 and BCL-xL, samples were fixed for 20 min on ice after surface marker and viability dye staining, followed by intracellular staining using BD Cytofix/Cytoperm staining kit (BD Biosciences) as per manufacturer’s specifications. For the detection of STAT3 and phospho STAT3, samples were fixed for 15 min at 37°C after surface marker and viability dye staining using Fixation Buffer (Biolegend), followed by permeabilization for 1h at −20°C with True-Phos™ Perm Buffer (Biolegend) as per manufacturer’s specifications. For the detection of IL-6, cells were cultured in the presence of brefeldin A (Sigma) for 4h, followed by fixation for 15 min at room temperature in 4% paraformaldehyde (Thermo Fisher). Samples were acquired on an LSR-Fortessa (BD Biosciences) with FACS-Diva software (BD Biosciences) and data were analyzed with FlowJo software (v10.3, Tree Star).

### Gene expression analysis

For gene expression profiling of C-IM and C-IM-IL6KO tumors, reporter positive cells (GFP+ hCD2+) were FACS sorted using a FACSAria III or a FACSAria Fusion (BD Biosciences). For gene expression profiling of in vitro derived activated B cells and plasmablasts, cells were FACS sorted at day 3 of LPS stimulation according to the expression of CD19 and CD138 markers (activated B cells – CD19^+^ CD138^neg^; plasmablasts – CD19^low^ CD138^+^). RNA was extracted using AllPrep DNA/RNA Mini and Micro Kits (Qiagen) as per manufacturer’s specifications. RNA sequencing was performed at The Francis Crick Institute Advanced Sequencing Unit. RNA sequencing was carried out on the Illumina HiSeq 2500 and 4000 platforms and typically generated around 25 million 101bp strand-specific paired-end reads per sample. Adapter trimming was performed with cutadapt (version 1.9.1) (Martin M, 2011) with parameters “--minimum-length=25 --quality-cutoff=20 -a AGATCGGAAGAGC –A AGATCGGAAGAGC”. The RSEM package (version 1.3.0) (Li and Dewey, 2011) in conjunction with the STAR alignment algorithm (version 2.5.2a) (Dobin et al., 2013) was used for the mapping and subsequent gene-level counting of the sequenced reads with respect to mm10 Ensembl genes downloaded from the UCSC Table Browser (Karolchik et al., 2004) on 19th February 2016. The parameters used were “--star-output-genome-bam –paired-end --forward-prob 0”. Differential expression analysis was performed with the DESeq2 package (version 1.12.3) (Love et al., 2014) within the R programming environment (version 3.3.1) (REF). An adjusted p-value of <= 0.05 was used as the significance threshold for the identification of differentially expressed genes. The Subjunc aligner from the Subread package (version 1.5.1) (Liao et al., 2013) was used for the quantification of Ighv transcripts as described in Shi et al., 2015. The parameters used were “--allJunctions -I 16 -u”. Gene set enrichment analysis (GSEA) (version 2.2.3) (Subramanian et al., 2005) pre-ranked analysis was performed using the Wald statistic with respect to custom signatures obtained from the literature. All parameters were kept as default except for enrichment statistic (classic), min size (5) and max size (50 000). Gene set enrichment analysis for differentially expressed genes was performed by Gene Ontology Pathway and Biological processes using GeneGo MetaCore (https://portal.genego.com/).

### Quantitative real time PCR

Total RNA was reverse transcribed using the SuperScript® III First-Strand Synthesis System with Oligo(dT)_20_ (Invitrogen). For qRT-PCR analysis, the Power SYBR Green Master Mix was used, followed by quantification with the StepOnePlus System (Applied Biosystems). Samples were assayed in duplicate, and messenger abundance was normalized to that of HPRT1. Primers sequences used for Prdm1, Xbp1, Sdc1 and Irf4 amplification can be found in the table attached.

### Statistical analyses

Data were analyzed using unpaired two-tailed Mann-Whitney test for two-way comparisons. Statistical significance for multiple comparisons was calculated using the False Discovery Rate approach by using the Two-Stage Step-Up method of Benjamini, Krieger and Yekutieli. P-value of 0.05 or less was considered significant. Prism (v8, GraphPad) was used for statistical analysis. Data in text and figures are represented as box plots with floating bars representing min and max values, with median value represented as a line.

### Analysis of tumor clonality

Southern blotting of EcoRI-digested genomic DNA from C-IM tumors using a JH probe spanning the JH4 exon and part of the downstream intronic sequence.

### Serum protein electrophoresis

Serum from C-IM tumor mice and C-EYFP aged mice was diluted 1:2 in barbital buffer and analyzed on a Hydragel K20 system (Sebia) according to manufacturer’s instruction.

